# Mechanism of sensor kinase CitA transmembrane signaling

**DOI:** 10.1101/2023.02.06.527302

**Authors:** Xizhou Cecily Zhang, Kai Xue, Michele Salvi, Benjamin Schomburg, Jonas Mehrens, Karin Giller, Marius Stopp, Siegfried Weisenburger, Daniel Böning, Vahid Sandoghdar, Gottfried Unden, Stefan Becker, Loren B. Andreas, Christian Griesinger

## Abstract

Membrane bound histidine kinases (HKs) are ubiquitous sensors of extracellular stimuli in bacteria. Here, we used solid-state NMR in conjunction with crystallography, solution NMR and distance measurements to investigate the transmembrane signaling mechanism of a paradigmatic citrate sensing membrane embedded HK, CitA. Citrate binding in the sensory extracytoplasmic PAS domain (PASp) causes the linker to transmembrane helix 2 (TM2) to adopt a helical conformation. This triggers a piston-like pulling of TM2 and a quaternary structure rearrangement in the cytosolic PAS domain (PASc). Crystal structures of PASc reveal both anti-parallel and parallel dimer conformations. An anti-parallel to parallel transition upon citrate binding agrees with interdimer distances measured in the lipid embedded protein using a site-specific ^19^F label in PASc. These data show how Angstrom scale structural changes in the sensor domain are transmitted across the membrane to be converted and amplified into a nm scale shift in the linker to the phosphorylation subdomain of the kinase.

**One-Sentence Summary:** Transmembrane signal transduction of a PAS-domain containing histidine kinase occurs via a piston-like pulling of a transmembrane helix, and amplification by cytoplasmic PAS domain dimer rearrangement.

## Main Text

In microorganism, membrane receptors are essential for processing extracellular stimuli. As a part of the ubiquitous two-component signaling systems (TCS), membrane-bound sensor histidine kinases (HKs) are one of the most abundant classes of prokaryotic membrane receptors. Many bacteria contain dozens to hundreds of sensors for perceiving various environmental stimuli (*1*). The transmembrane signaling pathway in extra-cellular sensing TCSs follows a common scheme: the initial signal switching the HK from the ligand-free state to the ligand-bound state is generated by cue perception in the sensor domain, which, through a transmembrane (TM) domain and cytoplasmic **P**er-**A**rnt-**S**im (PAS) or HAMP (named after the proteins **H**istidine kinases, **A**denylate cyclases, **M**ethyl accepting proteins and **P**hosphatases) domains, leads to cross-phosphorylation in the dimeric kinase core at a histidine residue (DHp). Subsequently, the kinase domain phosphorylates the respective response regulator (RR) protein, resulting ultimately in the regulation of target gene(s). Transmembrane (TM) signaling mechanisms were postulated, based on the structures of the isolated extracytoplasmic receiver domain for KinB (*2*), TorT/TorS (*3*), LuxPQ (*4*), CitA (*5, 6*), NarX (*7*), and full-length DcuS (*8, 9*). The structure of the complete cytoplasmic region has been solved for some proteins, such as VicK, YF1 and HK853 (*10–12*). However, direct structural insight into the transmembrane signaling process is needed for PASc containing HKs.

The citrate sensor HK (CitA) is an attractive HK to study, because it is functional without any auxiliary proteins (*13*). It is a member of the PAS domain containing HKs which comprise 30 % of all HKs (*14*). CitA belongs to a sensor kinase subfamily (*15*) of HKs, which can bind to citrate or C_4_-dicarboxylates. This class of sensor HK has two PAS domains, a periplasmic (or extracytoplasmic) ligand binding domain (PASp) and a cytoplasmic transmitter domain (PASc). The PASp and PASc domains are connected to the TM2 helix by two linkers: the PASp/TM2-linker and the TM2/PASc-linker (topology and domain organization shown in Figure 1A and 1B). The PASp domain alone in both the citrate bound and the citrate free states has been studied by crystallography and solution state NMR (*6*). However, structural changes in PASc which would reveal the transmembrane signaling mechanism remain uncharacterized. Previously, we reported a piston-like motion in the PASp domain along with dynamics changes in the PASc domain of CitA from the thermophilic bacterium *Geobacillus thermodenitrificans* (GT) upon switching from the free to the bound state by citrate binding (*6*). PAS-containing crystal structures suggest a variability of structural arrangement between the PAS core and its N-terminal helix (*16*), including the observation of both a parallel and a 120° rotated dimer in one of the cytosolic PAS domains in the HK KinA (*17*). In addition, mutagenesis studies have also provided evidence for dimer rearrangement (*16*) upon the signaling event. Here, we report structural changes in the TM2 helix and its associated linkers at single-residue resolution, by using chemical shift information from ^1^H-detected solid-state NMR spectra. We used a bilayer-embedded CitA construct, referred to as the CitApc construct, that contains all essential elements for transmembrane signaling (PASp, TM helices and PASc) and the fully functional R93A mutant (*6*). With this construct, we detected a nanometer scale dimer rearrangement of the cytosolic PAS domains that occurs with the addition of citrate.

**Fig. 1.**
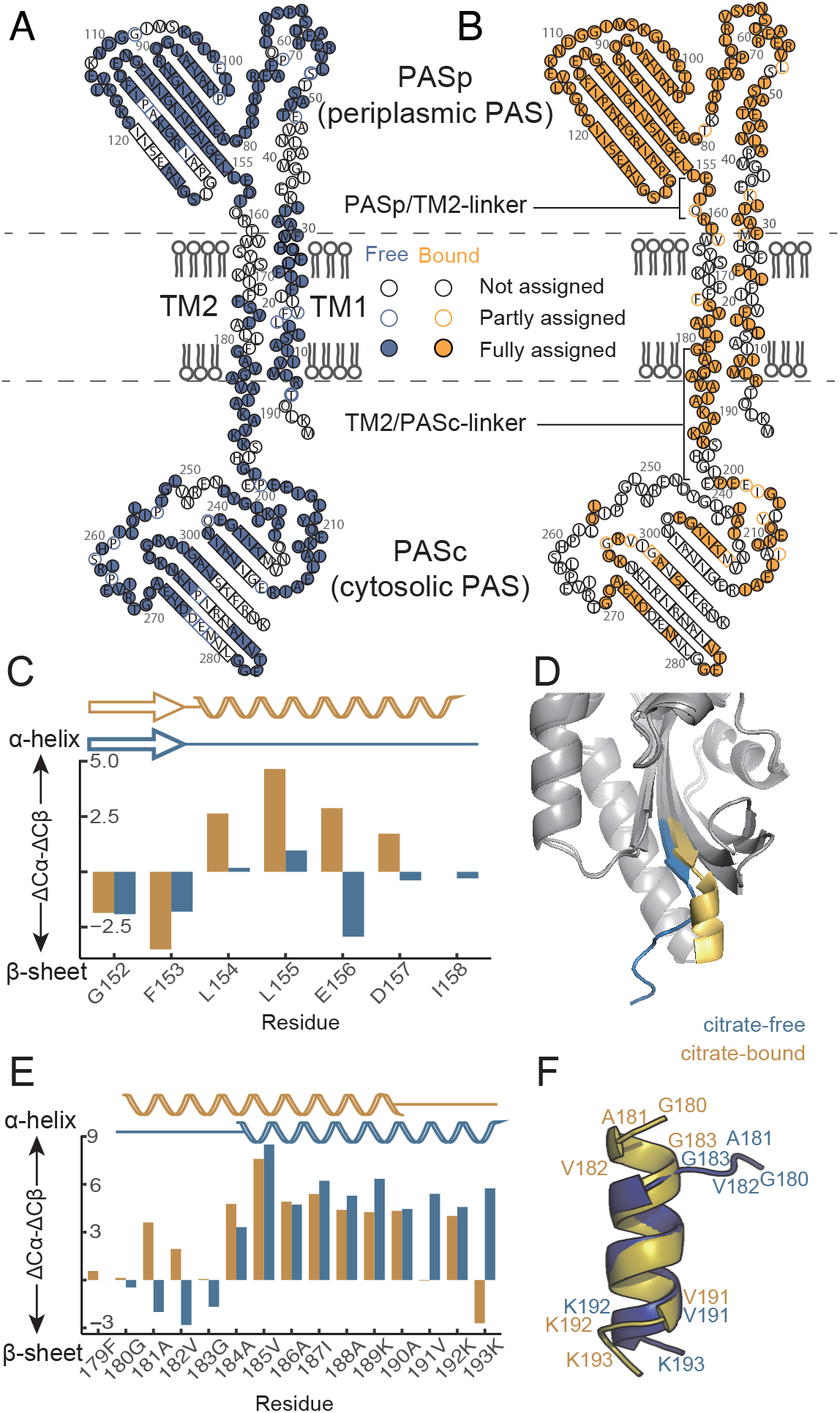
Sequence specific assignment with ^1^H-detected MAS NMR revealed secondary structural changes of CitA upon citrate binding. (A) Assigned residues shown in the topology map in the citrate bound and (B) the free state of the CitApc construct. (C) Citrate binding causes chemical shift changes in the PASp/TM2-linker. The contraction of the overall *β*-scaffold of PASp leads to a random coil to *α*-helix transition in the PASp/TM2-Linker. (D) Superposition of the citrate-bound GT PASp domain structure (PDB: 8BGB) with the citrate-free structure (PDB: 8BIY, SI tables 9, 10, 12 and 13). The border of this change ranges from L154 (previously described in Salvi *et al. (6)*) to I158. (E) Secondary chemical shift changes in the TM2/PASc-linker. (F) The structural model calculated using torsion angle restraints predicted with TALOS-N visualizes the N-terminal contraction and C-terminal loosening of TM2/PASc-linker upon citrate binding.

The extracytoplasmic helix (P-helix) at the C-terminal end of the receiver domain (PASp) is a consensus structural feature of HKs (*18*) and has been confirmed previously with ^13^C-detected solid-state NMR for the residue L154 in the ligand-bound state (*6*). Based on chemical shift assignment offered by proton-detected solid-state NMR experiments (Figure 1A and B, BMRB ID: 51759 and 51760), we are able to extend this helical characterization to four more amino acids up to I158, which is only one residue away from the predicted membrane embedded region based on cysteine accessibility in the homologous protein DcuS (*8*) (Figure 1C). This result is in agreement with the conversion of the disordered region downstream of the last β-strand in the PASp core (residues 152-158) in the citrate-free state to a helix (P-helix) extending to the TM2 helix, in the bound state (Figure 1D). This helix formation supports the proposed piston-like upward motion of the TM2 helix upon citrate binding (*6*).

Chemical shift assignments of the linker between the TM2 and PASc (*19*) in CitApc also indicate changes in secondary structure upon addition of citrate (example assignment spectra for this region are shown in Figure S1-S3). Details of the assignment process based on a series of 3D spectra are included in the SI materials and methods. With the upward piston-like motion of the TM2 helix, ^180^GAVG^183^ at the N-terminus of the TM2/PASc-linker undergoes a transition from mostly extended (β-sheet-like) secondary chemical shifts to *α*-helical secondary chemical shifts in the bound state. This region also experiences a change in its solvent environment from water to lipid as observed by solvent transfer experiments (*20*) (Figures S4 and S5). Since both glycine and alanine have a higher tendency to form an *α*-helix in a hydrophobic environment (*21*), the upward shift of the ^180^GAVG^183^ region into the membrane facilitates its contraction into a helical conformation. At the C-terminus of the TM2/PASc-linker, the ^192^KK^193^ motif does the opposite and loosens into a more extended structure when citrate is bound. This is seen from a change in secondary chemical shift from positive to negative for residues K192 and K193 (Figure 1E) and visualized in a structural model calculated based on torsion angle restraints predicted by TALOS-N (*22*) (Figure 1F). Cysteine cross-linking performed in DcuS TM2 and the linker also showed a reduced dimerization of the linker region in the presence of the ligand (*9*), which is compatible with a more extended conformation of the linker. Furthermore, a loosened ^192^KK^193^ linker would explain the faster dynamics in the PASc domain in the citrate-bound state indicated by the lower visibility of the PASc domain in cross-polarization (CP) based solid-state NMR spectra of CitApc with both ^13^C and ^1^H detection (*6*) (Figure 1A and 1B). This internal dynamics of the PASc domain limits the amount of assignable residues for unambiguous structure calculation of the PASc domain in CitApc by NMR (Figure 1A and B). Therefore, we applied NMR-based distance measurements guided by our crystal structures of the PASc domain (*23, vide infra*).

We observed more than one quaternary structure for the soluble PASc dimer in both crystals and in solution, in line with biochemical evidence suggesting that modulation of the PASc dimer interface is involved in the signaling process (*24–26*), We first generated functional mutants in the PASc domain from *G. thermodenitrificans* CitA, based on functional mutation *loci* in *E. coli* DcuS (*26*) identified through systematic mutagenesis. Function of the mutants was assessed by a β-galactosidase assay performed in *E. coli* CitA reporter strains (Figure 2F). The N288D mutation in the PASc domain retains the same function as the wild-type (WT) CitA. And, interestingly, the isolated N288D PASc domain shows an anti-parallel dimer arrangement in the crystal structure (Figure 2A, Table S11 and S14, PDB code: 8BJP), whose N-terminal helices run in opposite directions and, together with the N-terminus of the major loop of the PAS domain core (N245 to T252), form an anti-parallel dimer interface. In contrast, in the crystal structure of the WT PASc domain (*23*) (Figure 2B), the N-terminal helices of the dimer run in the same direction and form a parallel dimer interface. Notably, anti-parallel and parallel dimer forms are not domain-swapped dimers as seen in the cases of human βB2-crystallin (*27*) and cyanovirin-N protein (*28*), since they share the same monomer structure. The parallel dimer arrangement was previously reported in VicK (*11*) but the anti-parallel arrangement had not been reported. Both the WT PASc and N288D PASc eluted as a dimer from a size exclusion column (SEC) down to a concentration of 1 μM, maintaining the dimer state in solution as hypothesized for membrane bound CitA (Figure S6). In agreement, the parallel or anti-parallel dimer arrangements were found to be the dominant forms in solution, respectively, for the WT or N288D PASc domain based on the following two methods. First, the surface accessibility profile was measured from solvent paramagnetic relaxation enhancement (sPRE) and the measured intensity ratio profiles of the N-terminal helix distinguish between the dimer forms of WT (BMRB ID: 51764) and the N288D mutant (BMRB ID: 51765) PASc (Figure S7). Second, cryogenic optical localization in 3D (COLD) was used to measure the distance between two fluorophores covalently attached to the protein (*23*). The isolated PASc domains were labelled at the C-terminal residue 308 via introduction of an N to C mutation and attachment of the dye Atto647N. An inter-N308C distance of 41.3 Å was obtained in the N288D mutant PASc, which agrees with the expected distance in the anti-parallel dimer arrangement (Figure 2C and D). While some populations of dimers at smaller separations in Figure 2D cannot be excluded, the distribution of large distances about 41.3 Å d indicates that antiparallel arrangement dominates the PASc dimer. Regarding WT PASc, an inter-N308C distance of 9.0 Å is observed, which fits the expected distance based on the crystal structure of the parallel dimer (*23*).

**Fig. 2.**
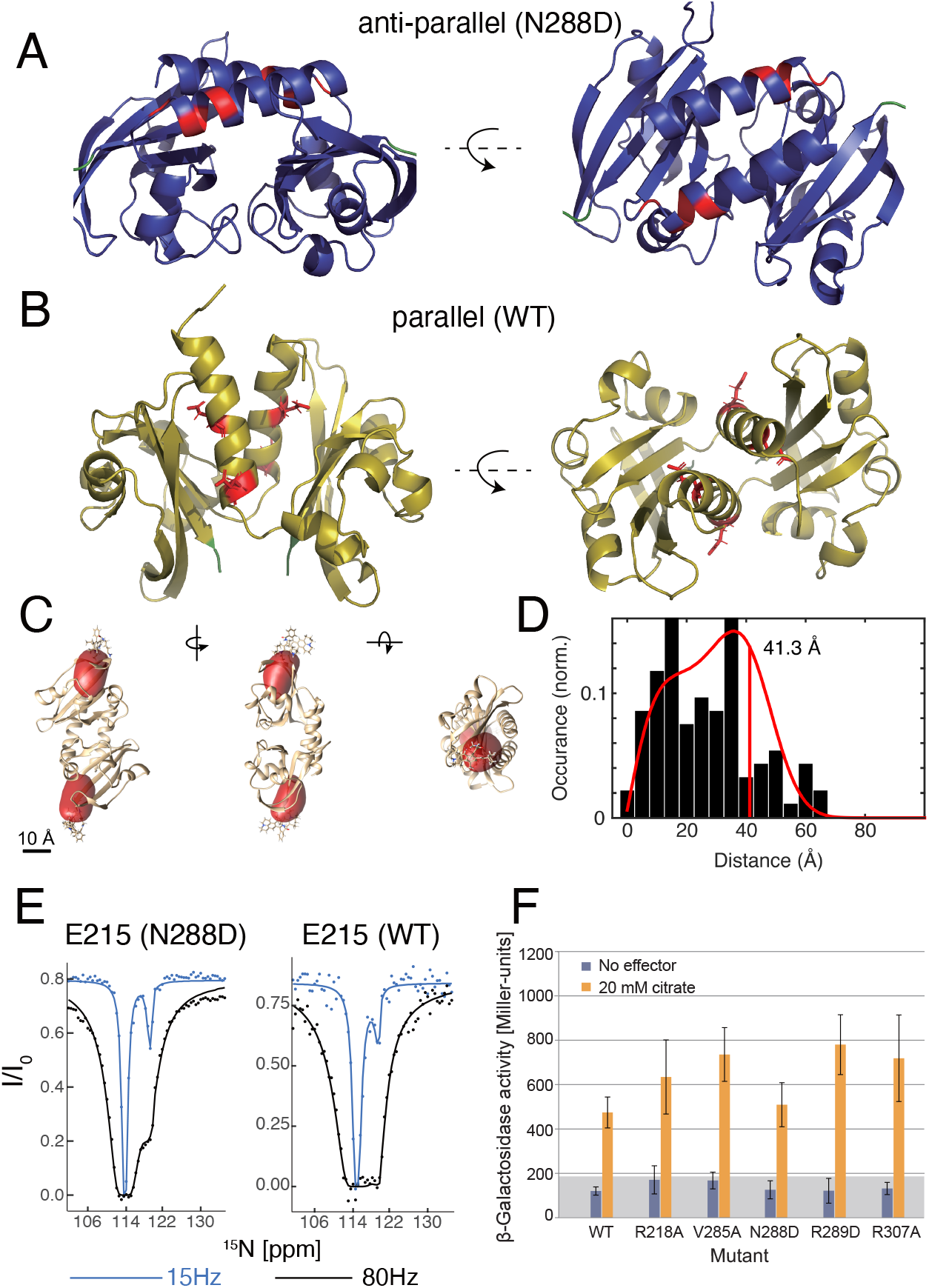
Residues at the dimer interface of isolated PASc show CEST exchange profiles, and COLD distance measurement confirms dimer forms in crystals. Residues showing CEST exchange (red) plotted on the crystal structure of (A) the N288D PASc mutant (PDB: 8BJP), which has the anti-parallel dimer arrangement and (B) the WT PASc (PDB: 5FQ1), which has the parallel dimer arrangement. The C-terminal residue N308 is shown in green. (C) Reconstructed volumes (red) of the fluorophores plotted at an isovalue of 0.68 and overlayed with the crystal structure of the N288D mutant PASc. Views from three orthogonal directions are shown. (D) Histogram of the measured projected distances and the expected distribution (red curve) as determined by a model fit. The red vertical line at 41.3 Å shows the resulting expectation value of the inter-fluorophore distance. (E) Example CEST profile and fitting of the residue E215. The fractional populations of the minor state in the WT and N288D mutant PASc domain are 3.8% and 3.3%, respectively, based on the fitted exchange rates (Table S7 and S8). (F) The N288D mutant CitA has the same activity and reactivity to citrate as the WT CitA according to a *β*-galactosidase assay performed in *E. coli* reporter strain.

Transition between conformations of the isolated PASc domain is observed by chemical exchange saturation transfer (CEST), which detects lowly populated states when they undergo millisecond (ms) timescale conformational exchange with the main population (*29*). For the PASc domain in CitA, this CEST effect was observed mainly in the residues from the N-terminal helices that form the dimer interface in both the anti-parallel and parallel arrangements (Figure 2E and S8). This indicates a dimer interface rearrangement of the PASc domain in isolation where the PASc domains switch most probably between the parallel and anti-parallel dimer forms. Although this dimer exchange seen in the WT PASc is functionally more relevant, its occurrence in the N288D PASc mutant helps to rationalize why this point mutation does not disturb the functionality of the full length CitA (Figure 2F). Still, connecting a PASc domain conformation with a specific citrate receptor state requires a membrane embedded construct.

For this purpose, we measured the characteristic dimer distance at residue 308 in the CitApc construct embedded in lipid bilayers (Figure 3D). The same mutation used previously in COLD measurements was introduced for site directed ^19^F labeling, and the method Center-band Only Detection of EXchange (CODEX) (*30*) combined with dynamic nuclear polarization (DNP) was applied. DNP was important to provide sufficient sensitivity to detect the fluorine signal from the singly CH_2_COCF_3_-tagged protein in the liposome embedded sample. Equally crucial is the combination of sensitivity enhancement provided by the three magnetically equivalent fluorine atoms in the CF_3_ group, and proton to fluorine cross polarization (CP) (Figure S10). An inter-CF_3_ group distance shorter than about 20 Å is measurable by fitting the CODEX decay curve, and a distance larger than 20 Å can be determined in the absence of decay within the 250 ms of CODEX. The CODEX experiment performed in both the free and bound state of the CitApc construct can then distinguish between the anti-parallel and parallel dimer in the PASc domain, where the former has an inter-N308C distance of 40 Å and the latter of 11 Å based on their crystal structures (Figure 3B and 3C) and the COLD measurement (Figure 2E and 2F). A decay in the CODEX signal to half of the reference signal was observed when citrate was bound (Figure 3A). Fitting of the decay curve resulted in a distance of 13.6 ± 3 Å, corresponding to the inter-N308C dimer distance expected in the parallel dimer within experimental error and tag flexibility. Nearly no CODEX decay was observed in the citrate free state (Figure 3A), indicating an inter-N308C dimer distance of larger than 20 Å, as expected in the anti-parallel dimer. The minor decay is caused by a 30% population of the bound state in the citrate free sample that was estimated from the intensity of the

**Fig. 3.**
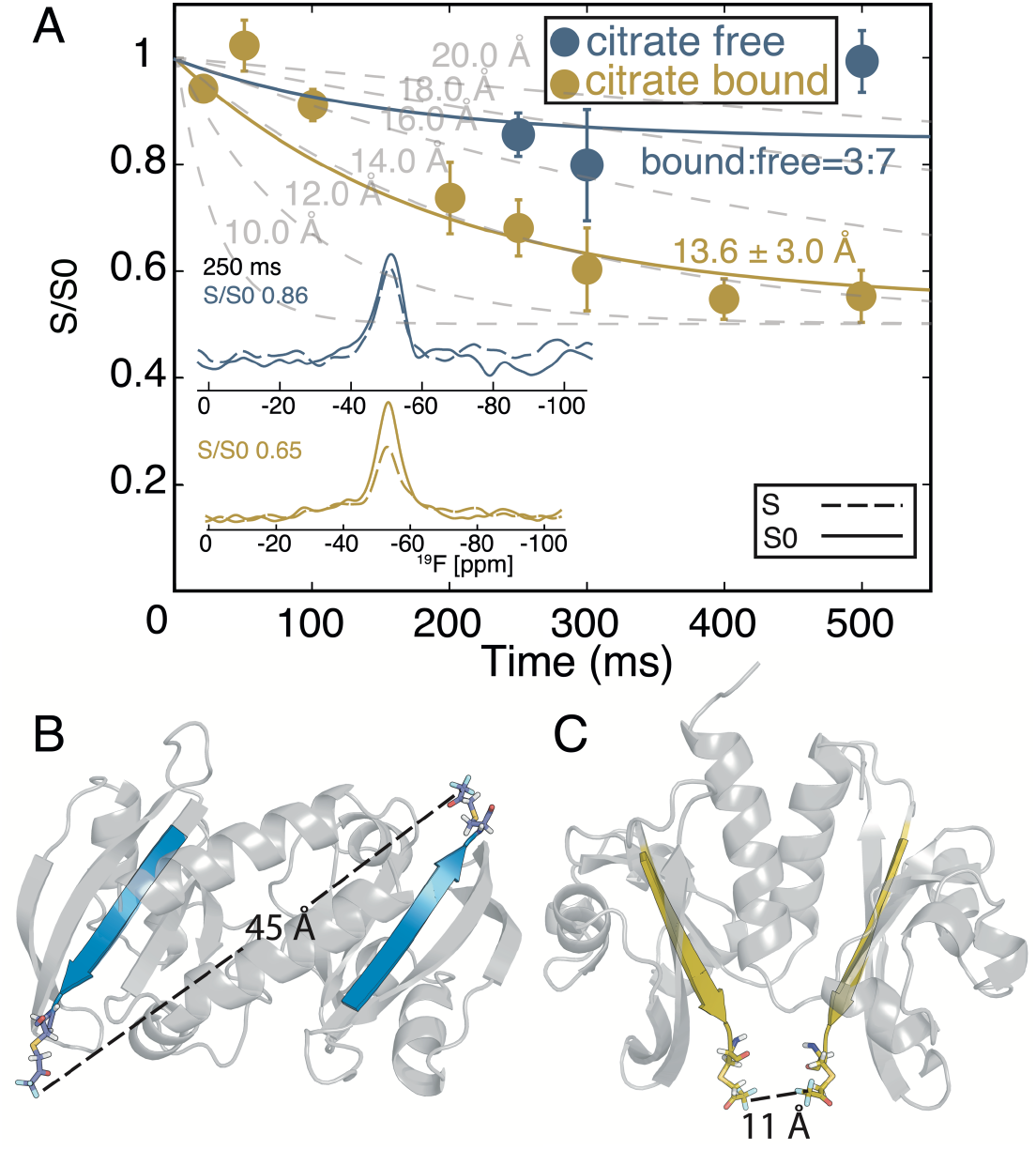
The C-terminal dimer distance of CH_2_COCF_3_ PASc observed by CODEX dephasing for both bound (yellow) and free (blue) states of the CitA CitApc. At the mixing time of 250 ms, the bound state CODEX signal decays to 0.65 of the reference experiment (A, inset yellow); the decay rate fits to an inter-CF_3_ distance of 13.6 ±3.0 Å, with the exponential decay fit curve of 0.5 * e^−0.0053*t^−0.5, matching the expected inter-dimer distance at the C-terminus of the parallel dimer (C). In contrast, the free state CODEX signal is almost the same as the reference experiment (A, inset blue) through all mixing times, corresponding to a large inter dimer distance at the C-terminus of more than 20 Å, agreeing with the anti-parallel dimer (B). Example CODEX decay curves at different inter fluorine distances are shown (A, grey). 30% of the bound state CitApc protein present in the citrate free sample caused the minor CODEX decay. The CODEX decay curve could be acquired beyond the ^19^F T_1_ of 321ms (Figure S10), thanks to an eight-fold DNP signal enhancement (Figure S9B).

H96 side chain peak in the (H)NH spectra characteristic of the bound form (Figure S13C). Considering this 30% population, a population weighted fit reveals a distance beyond 20 Å, and certainly larger than the inter-N308C distance found in the bound state. The dramatic change in the inter-dimer distance at the C-terminus of the PASc domain as identified with CODEX confirms a rearrangement in quaternary structure that is compatible with an anti-parallel to parallel dimer transition upon ligand binding. Importantly, CH_2_COCF_3_-tagged CitA is fully functional and responsive to citrate binding based on the ATP hydrolysis rate tracked using ^31^P NMR (*31*) (Figure S9). Upon citrate binding, a threefold increase in CitA’s ATP hydrolysis rate was observed, similar to the level of activity increase found using an *in vivo* assay (Figure 2D).

Our structural findings related to transmembrane signaling of the citrate binding sensor kinase CitA are presented in Figure 4. The large change in quaternary structure upon citrate binding detected by CODEX, is represented by the high-resolution structures of PASp and PASc dimers, and together with changes in secondary structure identified by NMR chemical shifts, the mechanism of transmembrane signaling is revealed. Structural changes include the P-helix formation, which promotes the piston-like pulling of the TM2 helix, and the amplification of this upward motion by the helical conversion of residues ^180^GAVG^183^, which border the membrane and the cytosol. This contraction into a helical conformation can be explained by the change of solvent environment as these residues are pulled up from aqueous phase into the membrane, becoming part of TM2. This contraction of the TM2/PASc-linker is compensated only partly by loosening of the ^192^KK^193^ motif close to the N-terminus of PASc, leading to a change in the residue 308 dimer distance from more than 20 Å to 13.6 ±3.0 Å based on CODEX data. These distances are consistent with a transition from the anti-parallel to a parallel conformation captured in crystal structures of the isolated PASc domains. Substantial dimer reorganization is not without precedent, as a 120° rearrangement has been observed in an isolated PAS-A domain (*17*). A transition to a parallel dimer, which aligns the N-terminal helices in the citrate bound state is likely caused by the shortening of the TM2/PASc-linker.

**Fig. 4.**
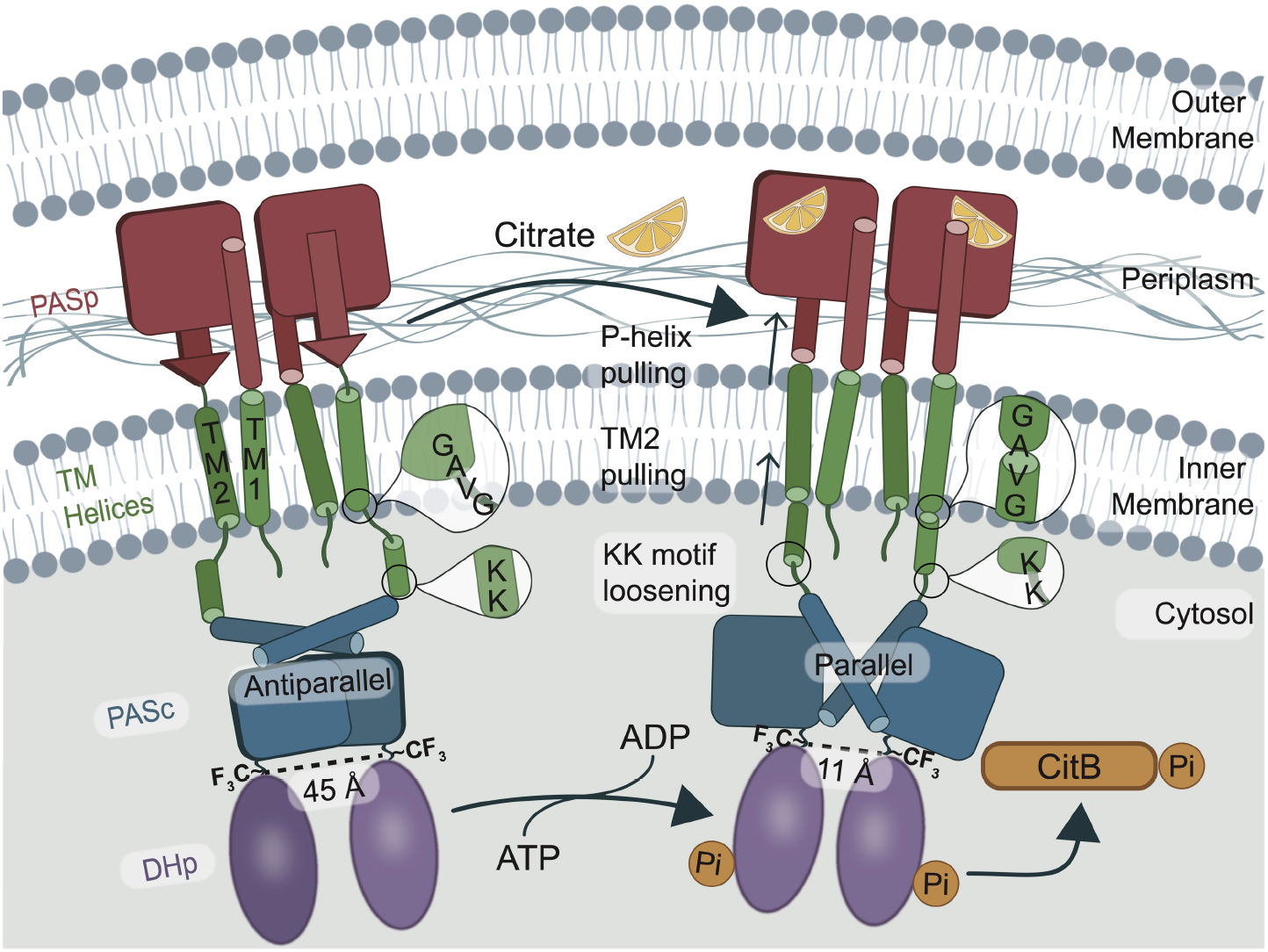
Mechanism of CitA activation by citrate, derived from a combination of structural studies of PASp, PASc and CitApc construct and activity assays of full length CitA. Important structural changes underlying the transmembrane signaling process of CitA include the P-helix formation, piston-like shift of the TM2 helix into the periplasm, an extension in the _192_KK_193_ motif, and the larger inter-dimer distance change at the PASc C-terminus. Altogether, the citrate binding has caused a large intracellular structural change, which is available to potentiate a difference in the CitA cross-phosphorylation and consequently, phosphorylation and activation of the RR CitB.

Several mechanistic details of CitA activation are similar to those observed in the HK NarQ, which has HAMP domains rather than PAS domains. NarQ uses HAMP as both its receptor and cytosolic connector domains. The crystal structure of NarQ without its kinase core revealed a piston-like motion for the TM helices upon activation similar to that detected in the PASp/TM2-linker and the TM2 helix of CitA. In contrast, the lever-like rearrangement of the NarQ HAMP domain was distinct from the structural rearrangement observed here in the PASc domain upon activation (*32*).

In summary, we have formulated a transmembrane signaling mechanism of CitA from the citrate binding at the PASp domain to the structural and dynamic changes in the PASc domain. Future studies will have to elucidate the arrangement of the PASc dimer in the context of the full-length CitA, and how its conformational dynamics affect the structure of the DHp dimer. Given the fact that a large number of HKs contain a cytosolic PAS domain, these findings of our study can be applied across a wide spectrum of bacterial HKs and also assist with the formulation of a uniform HK activation mechanism.

## Acknowledgments

We acknowledge discussions with Dariush Hinderberger on EPR results. We thank the beamline staff at SLS, X10SA for support with x-ray data collection.

## Author contributions

Conceptualization: CG, LBA, SB

Methodology: CG, LBA, SB, VS, GU

Investigation: XCZ, KZ, M. Salvi, KG, JM, SW, M. Stopp, BS

Funding acquisition: CG, LBA, VS, GU

Project administration: CG, LBA, SB

Supervision: CG, LBA, SB, VS, GU

Writing – original draft: XCZ

Writing – review & editing: all

## Data and materials availability

X-ray data of the GT PASp domain in citrate bound and citrate free forms, the GT PASc WT and N288D mutant are available at protein data bank (PDB), with the ascension numbers: 8BGB, 8BIY, 5FQ1, 8BJP. NMR assignment data of the GT CitApc in citrate bound and citrate free forms, the GT PASc WT and N288D mutant are available at biological magnetic resonance data bank (BMRB), with the ascension numbers: 51759, 51760, 51764, 51765.

## Supplementary Materials

Materials and Methods

Figs. S1 to S13

Tables S1 to S14

References (*33–60*)

