## Supplementary file for "Mechanism of sensor kinase CitA transmembrane signaling"

**This PDF file includes:**

Supplementary Materials and Methods
Figs. S1 to S13
Tables S1 to S14
References (33 to 60)

### Materials and Methods

#### Protein sample preparation

For producing approximately 100% deuterated, $^{15}N$, $^{13}C$-labeled C12A/R93A CitA1-309, CitApc, the plasmid coding for C12A/R93A CitApc was transformed into the strain C43 (DE3) (Imaxio, France). The R93A mutant was previously used in the isolated PASp domain to investigate its structure in both the citrate bound and citrate free form (*6*). Without the R93A mutation, the PASp domain in GT CitA binds the citrate molecule too tightly for a citrate free state to be produced. A single colony was transferred into a 2 ml minimal medium shaking culture and step-wise adapted to 100 % $D_2O$, followed by expression in minimal medium with 100 % $D_2O$, $^{15}N$-$NH_4Cl$ as nitrogen source and $^{13}C_6$-D-glucose as carbon source. For additional reverse labeling with isoleucine, valine, phenylalanine and leucine, 150 mg/L L-isoleucine-$d_{10}$, 100 mg/L L-valine-$d_8$, 50 mg/L L-phenylalanine-$d_8$ and 150 mg/L L-leucine-$d_{10}$ (Merck) were added to the expression culture shortly before induction. Purification was performed according to the established protocol (*6*). Briefly, after induction with IPTG, cells were incubated over night at 20 °C in a shaking culture before being harvested. The cell pellets were resuspended in TKMD buffer (50 mM Tris·HCl pH 7.0, 200 mM KCl, 5 mM $MgCl_2$, 5 mM β-mercaptoethanol, one spatula tip of DNAseI, 0.5 mM PMSF). The resuspended cells were lysed in a French pressure cell (20,000 psi). Cell debris was pelleted by centrifugation and supernatant spun down in an ultracentrifuge. The resulting pellet was re-suspended in Ni-NTA buffer (20 mM Tris·HCl pH 7.9, 500 mM NaCl, 10 mM imidazole, 5 mM β-mercaptoethanol, 0.5 mM PMSF, 4 % Triton X-100 (v/v)) and diluted using Ni-NTA-buffer without Triton X-100 to set the final detergent concentration to 0.8 % (v/v). The protein was purified using a 5 mL Ni-NTA column (GE Life Sciences). After washing, the detergent was changed on-column by washing with Ni-NTA-buffer supplemented with 1 % (v/v) laurydimethylamine-oxide (LDAO) instead of Triton X-100. For elution, the imidazole concentration was increased to 500 mM. Size exclusion chromatography (SEC) on Superdex 200 26/60 columns (GE Life Sciences) was performed as the final purification step using 20 mM Tris·HCl pH 7.4, 150 mM NaCl, 1 mM DTT, 0.3 % LDAO as running buffer. The purified C12A/R93A CitApc protein was reconstituted into 1,2-dimyristoyl-sn-glycero-3-phosphocholine (DMPC) and 1,2-dimyristoyl-sn-glycero-3-phosphatic acid (DMPA) liposomes (with DMPC to DMPA molar ratio of 9:1) at a lipid:protein ratio of 75:1 (mol/mol). The liposomes were taken up in 20 mM sodium phosphate, pH 6.5, 5 mM sodium citrate. The citrate free sample is made by incubating the citrate bound CitApc sample with the citrate free buffer (20 mM sodium phosphate pH 6.5) at 40 °C for a week. This process is monitored by tracking the intensity of the H96 side chain with 2D (H)NH spectra (Figure S13). The liposome samples were packed in a Bruker 1.3 mm rotor by ultracentrifugation.

Full length C12A/R93A/N308C GT CitA was expressed and purified using a pET16bTEV-vector based construct following the established protocol for the C12A/R93A/N308C GT CitApc construct with modifications. After purification by gel-filtration, the protein was loaded onto 4 ml $Ni^{2+}$-NTA resin and eluted with 20 mM Tris/HCl, pH 7.4, 150 mM NaCl, 500 mM Imidazole, 0.694 % (w/v) decylmaltoside (DM) and 0.5 mM Tris-(2-carboxyethyl)-phosphine hydrochloride (TCEP). After adding 6 mM ATP, the protein was reconstituted into DMPC liposomes at a lipid to protein ratio of 20/1 (w/w), removing the detergent with BioBeads (BioRad). The final concentration of the liposome-reconstituted protein in the buffer was adjusted to 100 μM.

The introduction of fluorine labels using cysteine alkylation by 3-bromo-1,1,1-trifluoroacetone (BTFA) is done according to the protocols published by the group of Schofield (*33–35*). Tagging of the C12A/R93A/N308C-GT CitA protein with BTFA was performed using 50 mM sodium phosphate, pH 7.0, 200 mM NaCl, 0.3% (w/v) LDAO in the gel-filtration step. Subsequently BTFA was added drop-wise to the protein solution as described for the BTFA-tagged CitApc, from a 100 mM stock solution in 50 mM sodium phosphate, pH 7.0, 200 mM NaCl. After incubation overnight on ice, the protein was loaded onto 4 ml $Ni^{2+}$-NTA resin to switch to 0.694 % DM and reconstituted in DMPC liposomes at a lipid to protein ratio of 20/1 (w/w), with 6 mM ATP added to the buffer. The final protein concentration was adjusted to 100 µM. To obtain the citrate bound state, buffer containing 5 mM sodium citrate was added to the sample and equilibrated overnight at 4 °C.

<u>sPRE measurement</u>

Solvent paramagnetic relaxation enhancement (sPRE) is measured and assessed according to published protocols (*36*, *37*). $^{15}$N WT and N288D PASc were doped with 2.5 mM of Gd-HP-DO3A, for measurement of the paramagnetic $^{15}$N-HSQC spectra. The intensity ratio between the paramagnetic and the diamagnetic (without Gd-HP-DO3A) $^{15}$N-HSQC spectra ($I_{sPRE}/I_0$) was used as the experimental sPRE value. A lower value is indication of higher solvent accessibility due to paramagnetic relaxation from the soluble gadolinium relaxation agent. All the spectra were recorded on a Bruker 400 MHz spectrometer equipped with a 5 mm triple channel room-temperature probe using an inter-scan delay of 2.5 s.

The predicted sPRE values from the crystal structures were obtained from the algorithm described by Öster et al. (*35*). The output of the algorithm is a number indicating the linear enhancement of the relaxation rate of a spin as a function of the paramagnetic agent concentration and high value corresponds to higher solvent exposure.

<u>CEST experiment</u>

$^{15}$N chemical exchange saturation transfer (CEST) data were recorded as described by Vallurupalli *et al.*(*29*). $^{15}$N radio-frequency field strengths of 20 and 80 Hz were applied together with $^{1}$H decoupling of 3.5 kHz during a relaxation delay of 400 ms. CEST data consisted of a series of 2D spectra, acquired in an interleaved fashion, corresponding to $^{15}$N irradiation offsets incremented in steps of 0.5 ppm. Additionally, a reference experiment for which the relaxation delay is set to 0 was recorded. CEST fitting was done using an in-house software described previously by Carneiro *et al.*(*38*).

<u>$^{31}$P NMR ATP hydrolysis assay</u>

Before the $^{31}$P NMR activity assay, approximately 150 $\mu$L full length CitAliposome sample was pelleted and then resuspended in 20 mM Tris/HCl pH 6.5, 50 mM KCl, 10 mM $MgCl_2$, 0.5 mM EGTA and 6 mM ATP, 0.5 mM Pefabloc (Roth, Germany), 0.02 % $NaN_3$. After the measurement was finished, the samples were centrifuged and the pellet was weighed to compare the amount of protein in the citrate bound and the citrate free sample. The final reaction rate normalized by the weight was compared.

The detection of ATP in solution, along with its degradation product, the inorganic phosphate, with NMR is straightforward, owing to the 100% natural abundance of the NMR-active nucleus $^{31}P$ and the large chemical shift separation (*39–41*) of the free phosphate. 1D phosphorus spectra were measured with a Bruker 400 MHz magnet. The spectra for full length un-tagged R93A/C12A/N308C CitA were measured with a room temperature QXI H/P-C/N/D five channel probe with a $B_1$ field corresponding to a 7.5 kHz nutation frequency. The spectra for $CF_3$-tagged R93A/C12AN/308C CitA were measured with a room temperature TBO H/F-C/N/P-D probe. All spectra were measured with an inter-scan delay of 1 s and an offset at -15 ppm. The number of scans was set to 1920. 10% $D_2O$ was used in all samples as lock signal. The measurement temperature for all CitA full length constructs was 37 °C. The initial speed of the reaction was used to measure the kinase activity since the only reactant (ATP) is added in large excess to the enzyme. ATP is stable under the experimental condition (Figure S12), so its auto-hydrolysis does not need to be taken into account. The error is estimated from the noise level of the spectra. According to zero-order kinetics in high excess, the consumption of the reactant in the complete reaction should follow a single exponential decay. At very short reaction time, the free phosphate (Pi) production approaches a linear equation with the slope corresponding to the ATP hydrolysis rate.

β-Galactosidase assay

β-Galactosidase as a reporter of the activity of CitA of *G. thermodenitrificans* was tested in the *E. coli* strain, IMW549 (*citA*$^-$ and *citC–lacZ*), that is deficient of *E. coli citA* but carries an *E. coli citC-lacZ* β-Galactosidase reporter fusion (*12)*. The strain was transformed with plasmids encoding CitA of *G. thermodenitrificans* or variants R218A, V285A, N288D, R289D, or R307A thereof, (pMW1652 and variants) and CitB *of E. coli* (pMW1653). The bacteria were grown without or with citrate (20 mM), and the *citC–lacZ* reporter gene activity (mean ± SD) was measured (*12)* in three biological replicates and four independent experiments.

Sequence specific assignment and statistics

We were able to acquire NMR spectra with high frequency MAS, homogeneous samples and deuteration in both the citrate bound and citrate free states (Figure S1). The sequence specific assignment process of the linker from residue A181 to K193 in citrate free and citrate bound state using the 3D spectra is shown in Figure S2. To further confirm the assignment in the TM helices, (which have a higher abundance of the residue types of I, V, F and L), we measured samples in which these residues were reverse labeled (without $^{13}C$ and $^{15}N$ labeling). Resonances belonging to I, V, F and L are expected to have no intensity in the spectra (Figure S3) from this reverse labeled sample, while in practice, incomplete elimination of valine signals was observed.

$^{1}H$-detected solid-state NMR spectra of CitApc, except the (H)N(CA)(CO)NH spectrum in the bound state, were measured at the field of 850 MHz with a Bruker 1.3 mm H/C/D/N four channel probe. The (H)N(CA)(CO)NH spectrum in the bound state was acquired at the field of 800 MHz with a Bruker 1.3 mm H/X/Y three channel probe. The peak lists extracted from the assignment experiments were used in automated assignment by FLYA (*42*), which aided in the initial process of sequence specific assignment. The lipid transfer H(H)NH spectra were not used for this purpose.

Spectra were processed with Topspin and assigned using Collaborative Computing for NMR (CCPN) analysis (*43*).

The sample temperature was set to 293 K adjusted with the chemical shift of water, using a cooling gas flow of 1350 L/h at 243K. Chemical shifts of $^{13}C$ are referenced relative to sodium trimethylsilylpropanesulfonate (DSS).The following 3D spectra have been acquired (*44–46*): (H)CANH, (H)CA(CO)NH, (H)(CA)CB(CA)NH, (H)(CA)CB(CA)(CO)NH, (H)CONH, (H)(CA)CO(CA)NH, (H)N(CO)(CA)NH. Additionally, 3D lipid transferred experiment H(H)NH (*20*) is acquired to confirm the assignment in the membrane contacting regions. Details of the experimental parameters are summarized in Table S3, S4, S5 and S6.

X-ray crystallography

All constructs were expressed and purified as described before for the WT PASc (*6*, *23*). The fragments comprising the PASp domain (amino acids 33-161) and the PASc domain (amino acids 200-309) of Gt CitA were cloned each into a modified pET28a vector. The R93A mutant of the PASp fragment and the N288D mutant of the PASc fragment were generated using a QuikChange II Site-Directed Mutagenesis Kit (Agilent). PASp wildtype and mutant protein were expressed in in *E. coli* strain BL21(DE3), the N288D mutant PASc protein was expressed in selenomethionine-labeled form in methionine-auxotrophic *E. coli* strain B834 (DE3) (Novagen). All three fragments were purified by immobilized metal affinity chromatography on Ni-NTA resin (Qiagen). The N-terminal His-tag was removed with Tobacco Etch Virus protease and final purification by size exclusion chromatography was performed on a Superdex 75 16/60 column (GE Healthcare). The purified WT and R93A mutant PASp proteins were dialyzed against 20 mM HEPES, pH 7.0, 50 mM NaCl. Protein concentration was adjusted to 20 mg/ml using a Vivascience 10 kDa MWCO concentrator. Finally, 1 mM sodium citrate was added to the WT Gt PASp sample. The N288D PASc protein was dialyzed against 20 mM sodium phosphate pH 6.5, 50 mM NaCl and the protein concentration was adjusted to 35 mg/ml.

Crystals of the three proteins were obtained by the vapor-diffusion technique with sitting drops. 100 nl of protein solution were mixed with 100 nl of well solution (1M tri-sodium citrate, 0.1M imidazole, pH 9.0 for WT PASp and 0.2M lithium sulfate, 0.9M sodium di-hydrogen phosphate, 0.6M di-potassium hydrogen phosphate, 0.1M N-cyclohexyl-3-aminopropanesulfonic acid (CAPS), pH 10.6 for the R93A mutant PASp; 0.4M magnesium chloride, 0.1M Tris/HCl, pH 8.5, 20.5% PEG 8000 for the N288D PASc mutant). Crystals grew within one week and were cryoprotected by transferring them for one minute to well solution supplemented with 30 % glycerol (WT PASp), 33 % glycerol (R93A PASp) and 20 % glycerol (N288D PASc) and flash-cooled by plunging them into liquid nitrogen. Diffraction data sets were collected at beamline X10SA, SLS, Switzerland (PILATUS 6M detector, Dectris). For PASp and its R93A mutant, only native data were measured at a single wavelength (see SI Tables S9 and S10). Three data sets (peak, inflection and high-energy remote, see SI Table S11) were collected for the N288D PASc. All data were processed with XDS (*47*) and scaled with SADABS (Bruker AXS). Space group determination and statistical analysis were done with XPREP (Bruker AXS).

The structure of the wildtype PASp domain was solved by molecular replacement with PHASER (*48*) using the crystal structure of the citrate bound *Klebsiella pneumoniae* PASp domain (*5*) (PDB code: 2J80) as search model. Refinement was performed initially with Refmac (*53*) alternating with manual model building in Coot(*49*). Final refinement was performed with phenix.refine (*50*). The structure of the R93A mutant was also solved by molecular replacement with PHASER (*48*) using the crystal structure of the WT Gt CitA PASp domain as search model. Refinement was performed as described for the WT domain. Refinement statistics for the WT and mutant Gt PASp domain structures are shown in SI Tables 12 and 13. The structure of the N288D PASc domain was solved by multi-wavelength anomalous diffraction. Normalized difference structure factors were calculated using SHELXC, followed by solution of the substructure with SHELXD. Phasing statistics are shown in SI Table 11. Phase extension, density modification and main chain auto-tracing was done with SHELXE (*51*). Ten cycles of automatic model building alternated with structure refinement by ARP/wARP (*52*) resulted in modeling of 98 % of the residues. Refinement was done by positional and B factor refinement in REFMAC5 (*53*) alternating with manual model building in COOT (*54*). Refinement statistics for the N288D PASc domain structure are shown in SI Table 14.

<u>COLD measurement and statistics</u>

The details of the COLD experimental setup, sample preparation and data analysis are described in SI Material and Methods published previously in Weisenburger *et. al.* (*23*). We recorded COLD measurements on a custom-built cryogenic microscope at liquid helium temperature (T = 4.3 K). Samples were prepared by spin-coating diluted PASc N288D proteins conjugated with Atto647N-maleimide (AttoTec GmbH) at N308C onto thoroughly cleaned UV-grade fused silica coverslips (ESCO). We estimated the protein concentration by UV/VIS spectroscopy to be about 15 µM using an extinction coefficient of $\epsilon_{280}$ = 38,400 $M^{-1}$ $cm^{-1}$. The labeling efficiency was determined to be about 65 %. Raw images of the fluorescent recordings were analyzed using custom-written software in MATLAB (The MathWorks). Localization images were generated and then fed into a single-particle reconstruction using the subspace EM algorithm (*55*). To compare our results with the protein structure, we attached Atto647N fluorophores to the N308C positions at the C-termini of the crystal structure (PDB ID: 8BJP) by modeling. We used the point symmetry of the volume maps of the 3D reconstruction and fitted them to the crystal structure using UCSF Chimera, which was also used for visualization. The resolution of the 3D reconstruction was estimated by Fourier shell correlation of two half data set reconstruction using Free FSC (IMAGIC). Comparison with the half-bit criterion yields a resolution of 1.9 Å. We have performed a total of 13 similar experiment sessions over a period of seven months. The published data include 2,133 identified proteins, which after filtering resulted in 188 proteins at a yield of 8.8 %. The filtering criterion was localization precisions better than 6.5 Å, as described in Table S2 from Weisenburger *et. al.* (*23*). We graphically display distributions by histograms for all relevant data in Figure 2F. For the 3D reconstruction and orientation fitting, datasets above a threshold localization precision of 6.5 Å were excluded.

<u>CODEX measurement under DNP</u>

All liposome samples are prepared for DNP experiments by adding TEMTriPol powder to the liposome sample until the solubility is saturated (around 20 mM). All samples are packed into 2.5 mm Phoenix MAS rotors and shock frozen by plunging into liquid nitrogen.

All CODEX experiments are measured with a previously published pulse program (*30*, *56*, *57*) (Figure S10) on a Bruker 600 MHz magnet, with a 2.5 mm Phoenix DNP probe equipped with

H/F double channel tuning at 20 kHz MAS and a set temperature of 90 K. CODEX is an NMR method based on spin diffusion and CSA recoupling to investigate slow dynamics (*30*) and oligomer number (*56*, *57*). The rate of spin diffusion is parameterized by a factor F(0). This empirically determined overlap integral (physically a calibration of the rate of spin diffusion at a particular MAS frequency and magnetic field) can then be used to fit distances in unknown systems. F(0) is the overlap of the single quantum lineshapes of the two spins(*56*):

$$F_{ij}(0) = \int f_i(\omega - \omega_i) * f_j(\omega - \omega_j) d_\omega \quad [1]$$

$\omega_i$ is the center of each peak, and $f_i$ and $f_j$ describe the intensity of the signal of each spin. The rate of spin diffusion is also affected by the dipolar coupling, $\omega_{ij}$, between two spins such that the spin diffusion rate between $i$ and $j$ ($k_{SD,ij}$) is:

$$K_{SD,ij} = 0.5\pi * \omega_{ij}^2 * F_{ij}(0) \quad [2]$$

The spin diffusion rate can be calculated from the CODEX curve using models described in literature (*49*, *56*, *58*), by the following equations:

$$M(t) = e^{-Kt} M(0) \quad [3]$$

Equation 3 describes the magnetization evolution, $M(t)$, with spin diffusion time (t). $K$ is the $n$-dimensional exchange matrix containing the rate constants $k_{ij}$ between exchange of two spins $i$ and $j$ (see eqn. 2), where $n$ is the total number of orientationally different spins in close enough vicinity to undergo magnetization exchange.

The spectral overlap function $F_{ij}(0)$ generally has substantial variation from sample to sample. Here, we took the previously published $F_{ij}(0)$ value of 30 to 60 $\mu$s, determined at a magnetic field of 14.1 T, under 20-40 KHz MAS for a variety of fluorine containing molecules (*59*). Nevertheless, since $\omega_{ij}$ depends on the inverse cube of the distance, changes in F(0) affect the determined distance according to the sixth root. This means that a two-fold range in F(0), e.g. 30-60 $\mu$s will only introduce about 12 percent error in the determined distances.

$$k_{ij} = 0.5\pi \cdot F_{ij}(0) \cdot \omega_{ij}^2 \quad [4]$$

Monoexponential fit of the CODEX curve was performed using the GNUPLOT fit function (*57*). A population weighted monoexponential function was used to model the CODEX decay observed in the citrate free CitApc sample with 30% population in the bound state, with the fitted spin diffusion rate $k$ of 0.0053 from the citrate bound CitApc sample.

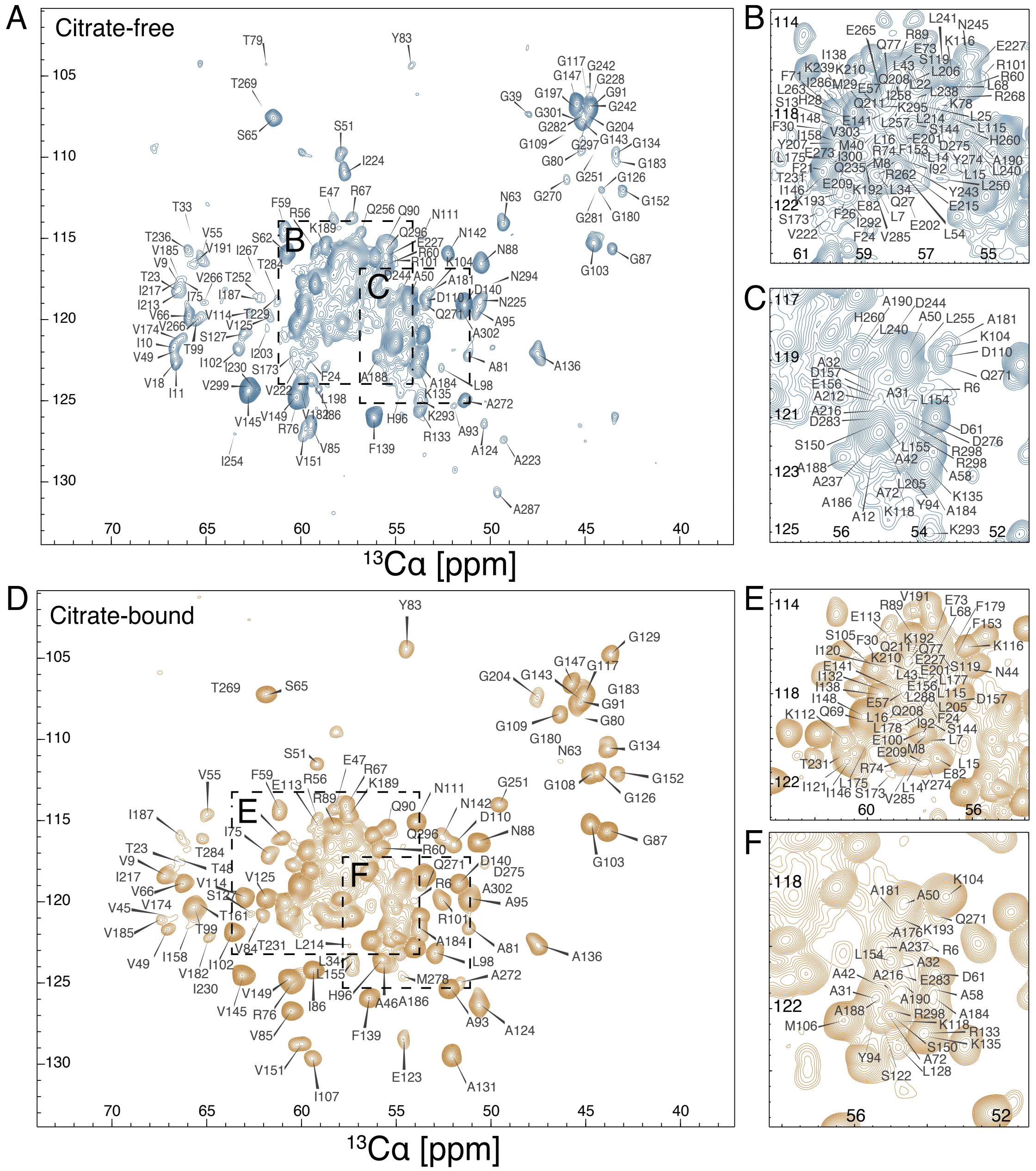


**Figure. S1. Spectra of bound and free lipid-embedded CitApc and residue specific assignment. (A to C) NC*α* projection of the (H)CANH spectrum in the free state, and (D to F) in the citrate bound state.** The assigned chemical shifts are mapped onto the spectra projections. The citrate bound state has more well-resolved peaks than the citrate free state. More assignments were found for the PASp domain in the citrate bound state, while the PASc domain assignments were more extensive in the citrate free state. All resonances from the assignment of the two state are included in Table S1 and S2.

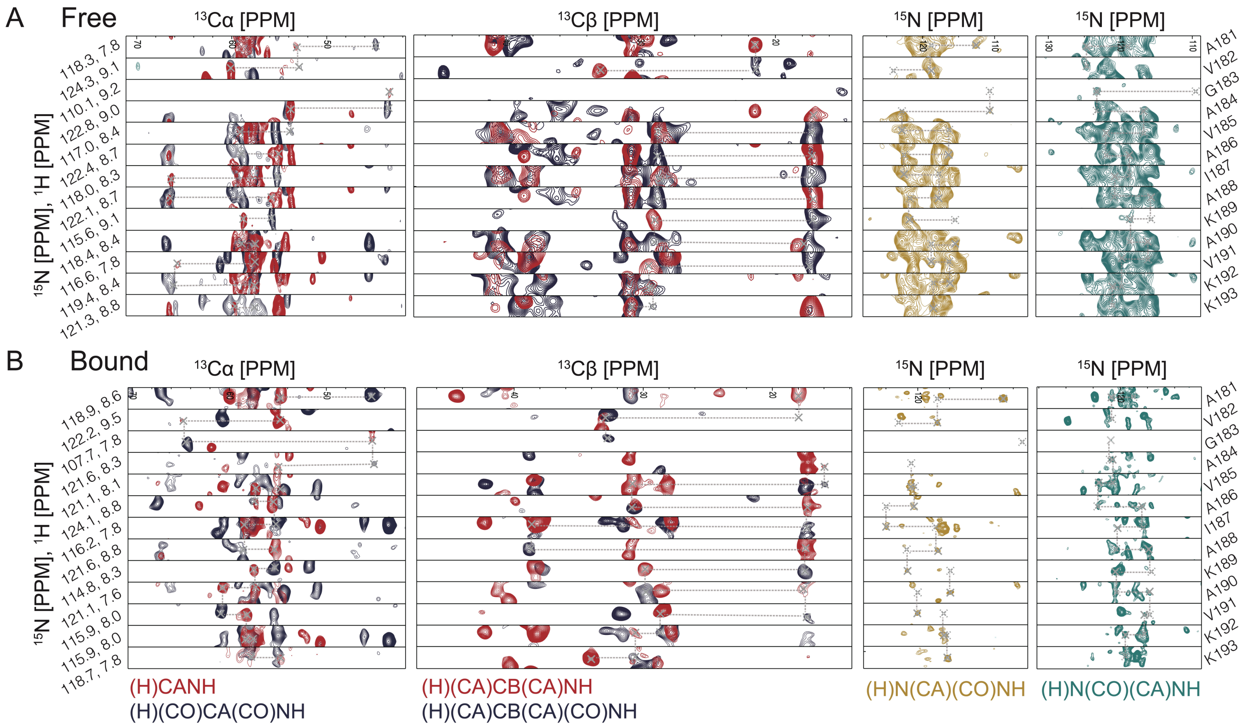


**Figure. S2. Sequence specific assignment of CitApc residues A181 to K193 in citrate free(A) and citrate bound (B) state.** The assignment process using (H)CANH, (H)(CO)CA(C))NH, (H)(CA)CB(CA)NH, (H)(CA)CB(CA)(CO)NH, (H)N(CA)(CO)NH and (H)N(CO)(CA)NH is shown. The same manual assignment process was done throughout CitApc in both states. The Cβ chemical shifts of K192 and K193 in the (H)(CA)CB(CA)NH spectra change to higher values with citrate binding. This indicates that the K192 and K193 residues have higher helix forming propensity in the citrate free state.

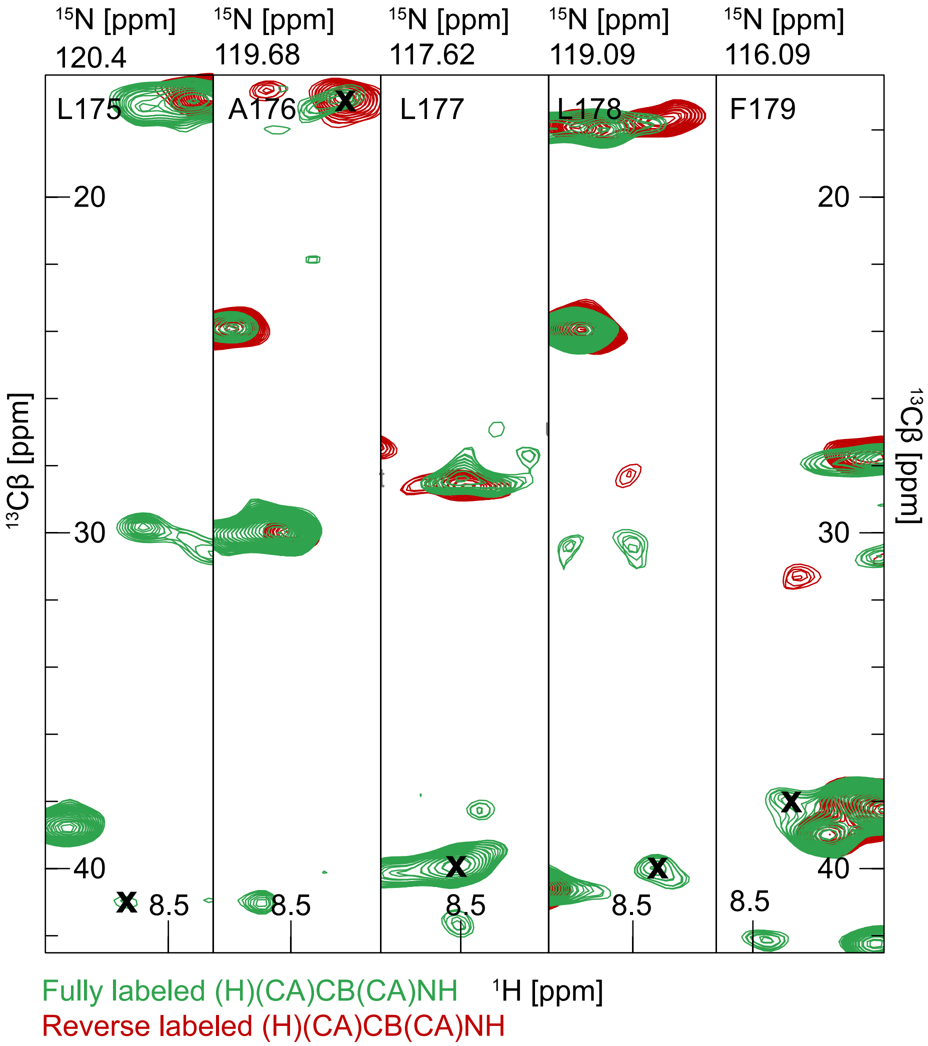


**Figure. S3. The IVFL reverse labeled CitApc sample helps with residue typing in the manual assignment process in particular in the TM helices.** An example is shown for the comparison of H(CA)CB(CA)NH spectra in both fully labeled (green) and reverse labeled (red) for residue L175 to F179 in the citrate bound state (peak position marked with bold x in each strip). The change in peak intensity along with the characteristic chemical shifts untangles the ambiguity of the assignment as the signal intensity for L175, L177, F179 disappeared, while the non-reverse labeled residue A176 remains the same.

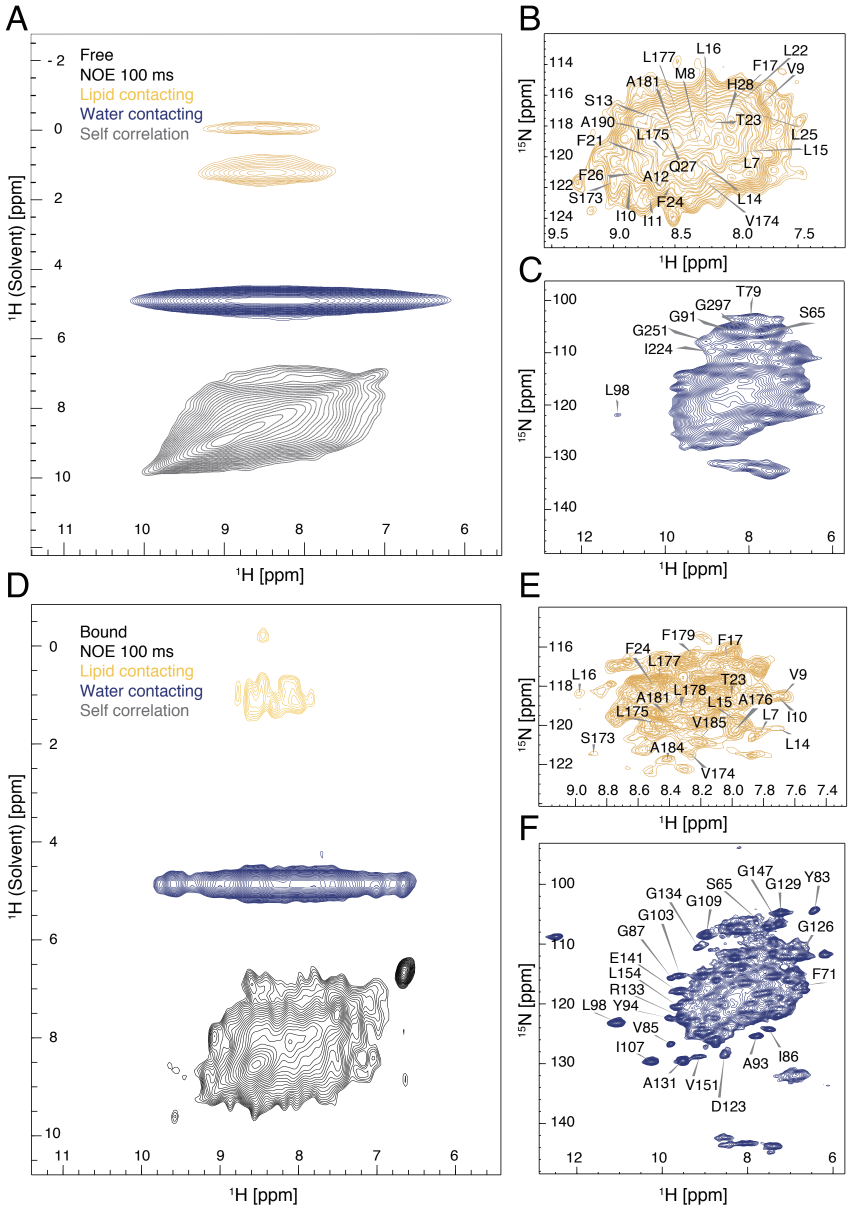


**Figure. S4. (H)HNH lipid contacting spectra of CitApc confirming chemical shift assignment of the TM helices in the citrate free (A, B, C) and citrate bound (D, E, F) state.** The additional proton dimension separates the signal from the proton and amide nitrogen dimension into water contacting (blue) and lipid contacting (yellow) parts (A, D). The protein-protein 'self-correlation' signals are shown in grey. The lipid tail group (0.7 ppm) contacting region in the free state has more intensity than in the bound state. The assigned TM helical resonances can be checked if they are contacting lipid on the H-N planes (B, E). The water (4.7 ppm) contacting region (C, F) in each state appears the same as the corresponding (H)NH spectrum since isolated peaks mostly come from either PASp or PASc, both of which have substantial exposure to water.

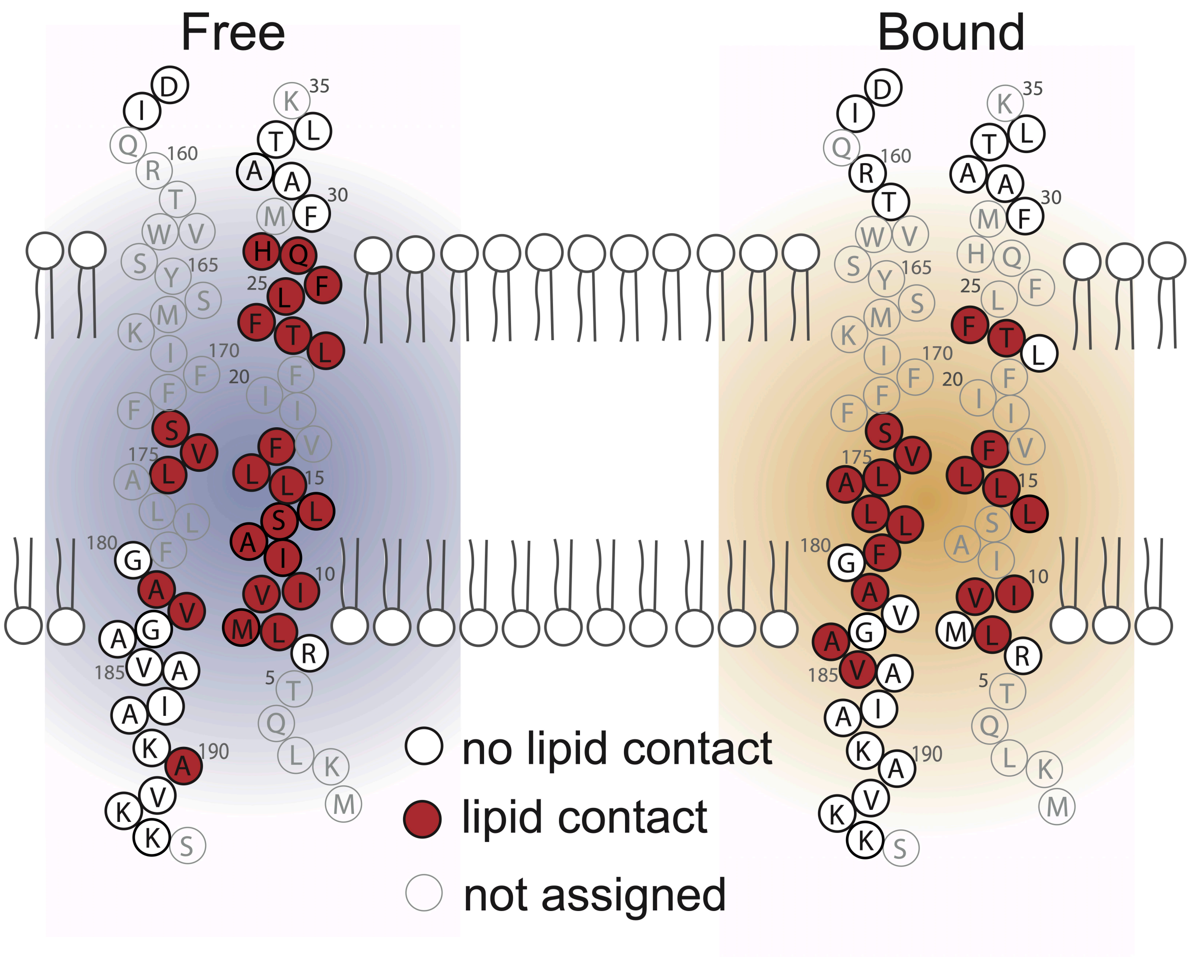


**Figure. S5. Lipid contacting residues in the TM helices mapped onto the CitA topology map in the free (left, blue) and bound (right, yellow) state of CitA.** Residues A184 and V185 show lipid contacts in the bound state but not in the free state. Residue A190 loses its lipid contact upon citrate binding. The border of lipid contact in the TM1 helix does not change while the border of lipid contact in the TM2 helix is shifted consistent with additional c-terminal residues entering the membrane.

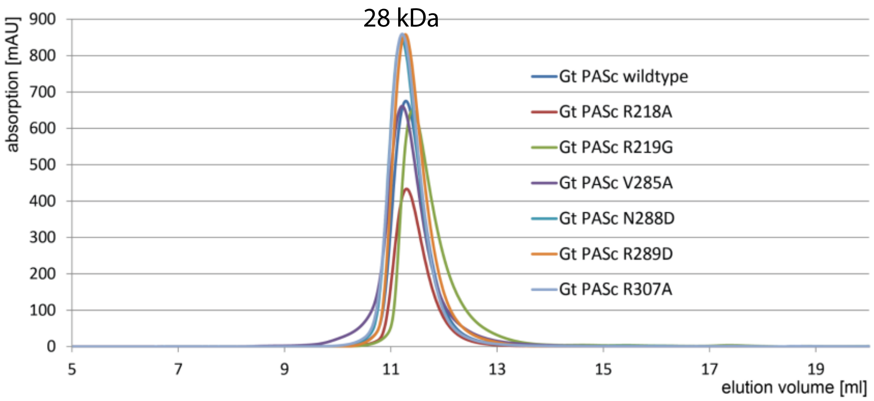


**Figure. S6. Size exclusion chromatography profile of *G.thermodenitrificans* CitA PASc mutants.**

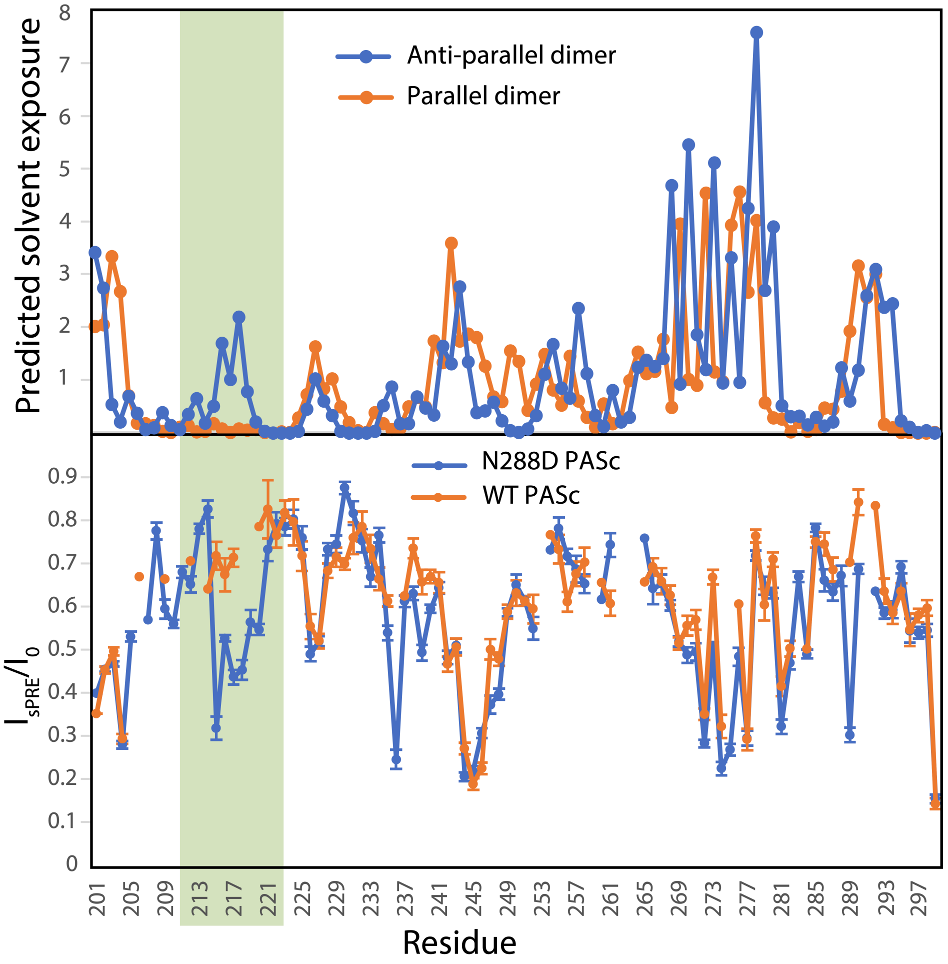


**Figure. S7. Predicted solvent exposure of the anti-parallel and parallel dimers from the crystals and the measured intensity ratios of the N288D mutant and WT PASc in solution.** (Top) Predicted solvent exposure of the anti-parallel dimer (blue) and parallel dimer (orange) from crystal structures of the N288D mutant and WT PASc. (Bottom) $^{15}$N-HSQC peak intensity ratios between the paramagnetic ($I_{sPRE}$) and diamagnetic ($I_0$) samples of the N288D mutant (blue) and WT (orange) PASc in solution. The sPRE profiles of two dimer forms are different from each other mainly at the N-terminal helix (green box). In this region, the anti-parallel dimer (top panel, blue curve) is predicted to have significantly higher solvent exposure, which corresponds to the lower intensity ratios in the N288D mutant PASc N-terminal helix in solution (bottom panel, blue curve).

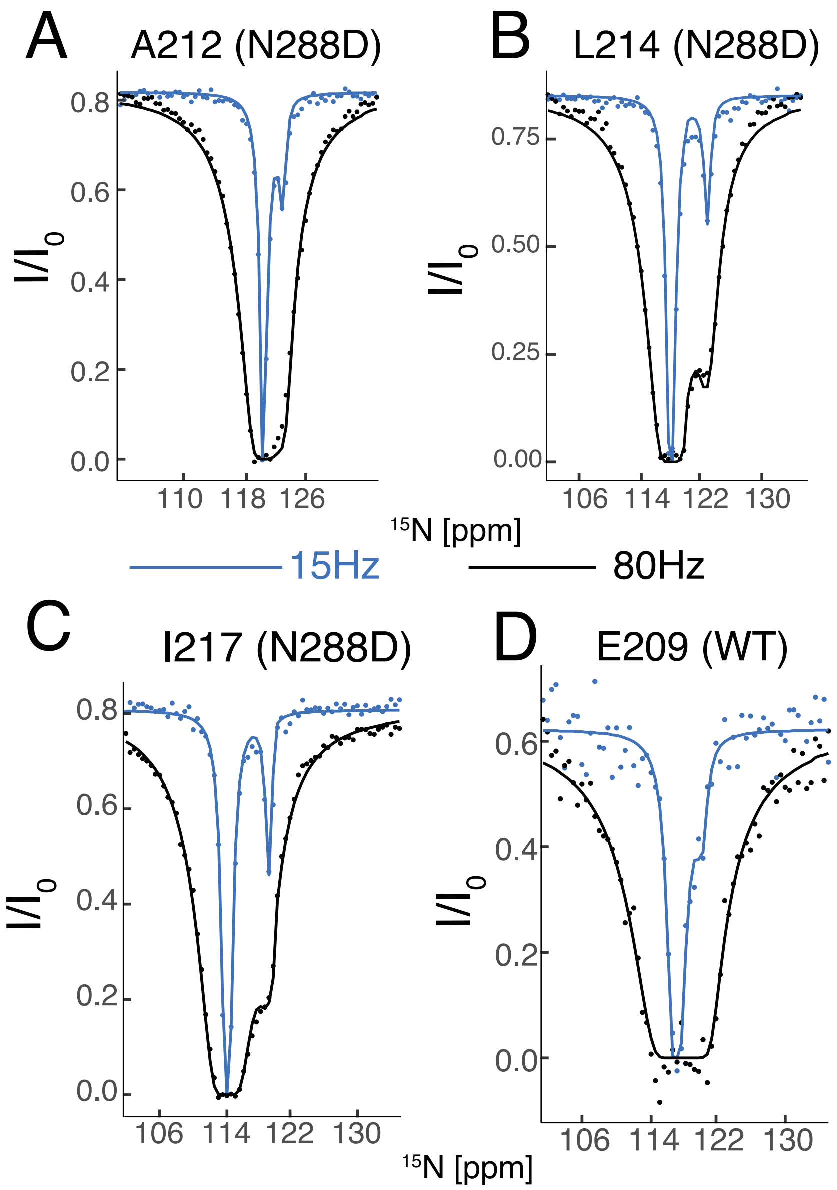


**Figure. S8. CEST profiles and fitting of residues (A) A212 in N288D PASc, (B) L214 in N288D PASc, (C) I217 in N288D PASc and (D) E209 in WT PASc.**

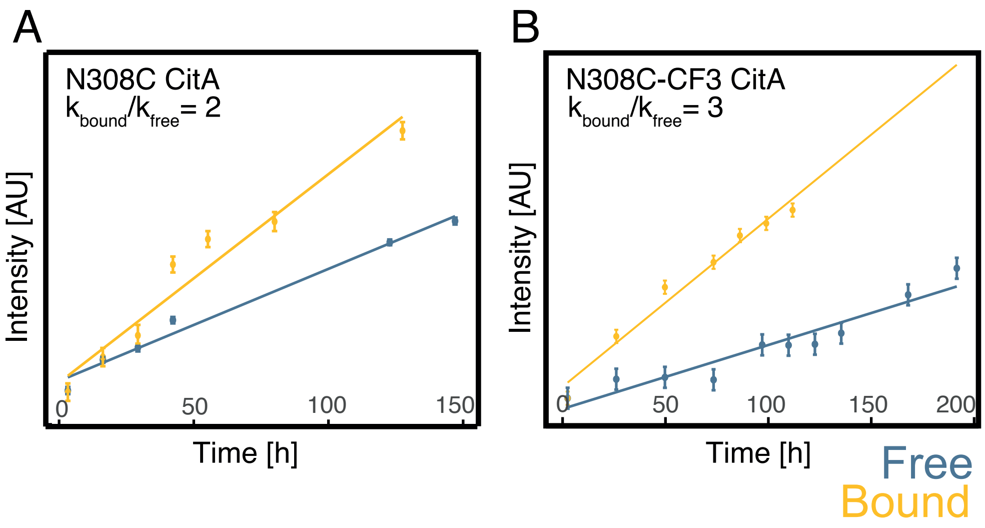


**Figure. S9. Kinase activity of full length C12A/R93A/N308C CitA without tag (A) and with $CF_3$ (B) from $^{31}P$ NMR assay.** The citrate bound state (yellow) in the non-tagged and $CF_3$-tagged CitA has two to three times higher activity compared with the citrate free state (blue).

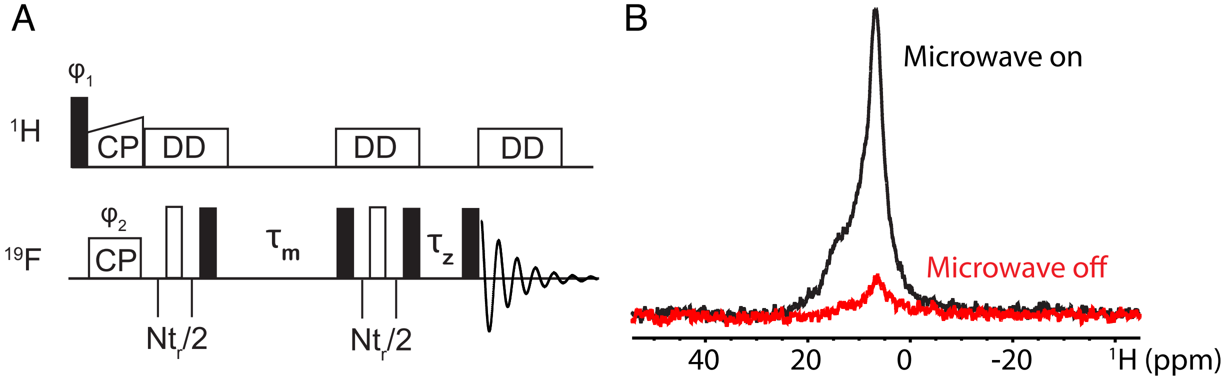


**Figure. S10. DNP enhanced CODEX measurement.** A) Pulse sequence for the HF CODEX measurement. $T_1$ compensation ($\boldsymbol{\tau_Z}$) and mixing time ($\boldsymbol{\tau_m}$) was set according to previous literature (*56*, *59*). A cross polarization (CP) contact time of 1.5 ms was used. For a proper recoupling of $^{19}F$ chemical shift anisotropy (CSA), N in the pulse sequence was set to 7. The sample temperature was 90 K, and the instrument a 600 MHz Bruker widebore spectrometer. The probe was a 2.5 mm Phoenix NMR (Loveland, CO) HFXY MAS DNP probe with MAS set to 20 kHz. B) a proton spectrum was used to characterized the signal enhancement from DNP. 8-fold enhancement was observed.

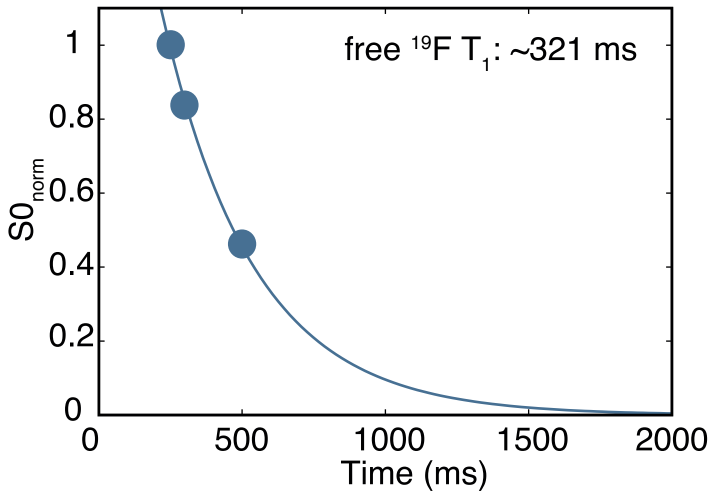


**Figure. S11. The $T_1$ of $^{19}F$ is 321 ms in the citrate free state, explaining the very weak signal observed with more than 500 ms of CODEX.**

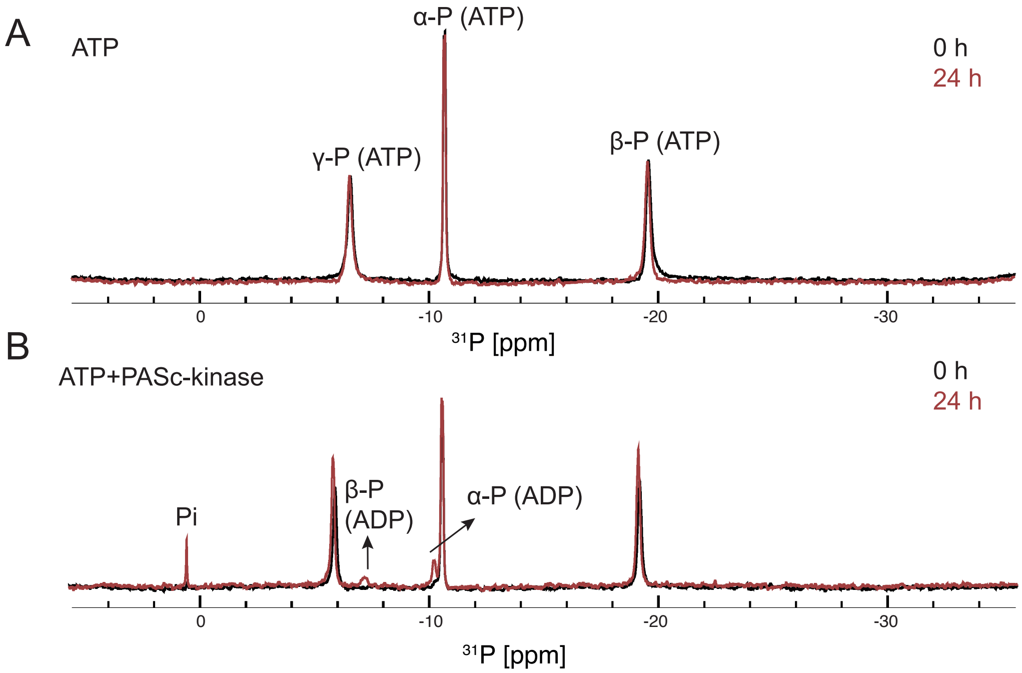


**Figure. S12. The addition of the PASc-kin WT construct hydrolyzes ATP into ADP (B), which is otherwise stable under the experimental conditions (A).** The 1D $^{31}$P spectrum of ATP does not change after 24 hours, indicating the stability of the ATP under the experimental conditions. The spectrum shows three peaks, corresponding to the signals of **β**-P, **α**-P and **γ**-P. The addition of PASc-kinase leads to the appearance of signals of the degradation products ADP (**β**-P and **α**-P in ADP) and free phosphate (Pi).

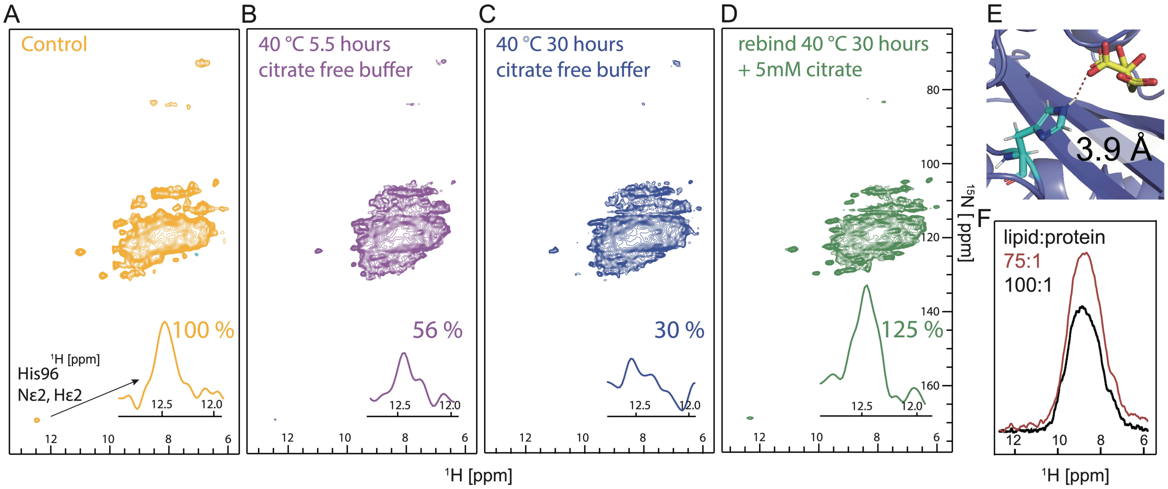


**Figure. S13. Citrate is removed from CitApc through incubation in citrate free buffer as tracked by the peak intensity of the H96 side chain.** (A) The citrate-bound spectrum. (B to C) the intensity drops to 30 % after 30 hours of incubation. (D) Addition of citrate to the buffer immediately restores the spectra to the bound state, indicating the reversibility of the protocol. (F) This citrate removal protocol optimizes sensitivity of the citrate free state, as it (red) increases the sensitivity by 30 % compared with the original protocol (black), which used a higher lipid to protein ratio of 100:1. (E) the hydrogen bonding interaction between H96 and citrate is shown as a dashed line. The loss of this hydrogen bond causes the H96 side chain to become unstructured and thus too dynamic to be detected by CP based (H)NH experiment.

**Table S1. Chemical shifts from $^{1}H$ -detected sequence specific assignment of the citrate bound state of CitApc.**

| Residue | H | N | C | Ca | Cb |
|---|---|---|---|---|---|
| 1Met | None | None | None | None | None |
| 2Lys | None | None | None | None | None |
| 3Leu | None | None | None | None | None |
| 4Gln | None | None | None | None | None |
| 5Thr | None | None | None | None | None |
| 6Arg | 9.36 | 120.09 | 173.9 | 54.1 | 27.89 |
| 7Leu | 7.92 | 120.01 | 177.77 | 58.13 | 40.94 |
| 8Met | 8.43 | 119.57 | 177.98 | 58 | 29.76 |
| 9Val | 7.72 | 118.45 | 176.88 | 67.2 | 29.03 |
| 10Ile | 7.73 | 118.42 | None | 60.01 | 40.44 |
| 11Ile | None | None | None | None | None |
| 12Ala | None | None | None | None | None |
| 13Ser | None | None | None | None | None |
| 14Leu | 7.71 | 120.52 | 178.6 | 57.9 | 40.45 |
| 15Leu | 7.92 | 120.17 | 177.11 | 56.29 | 41.24 |
| 16Leu | 8.95 | 118.51 | 178.24 | 58.76 | 40.15 |
| 17Phe | 8.06 | 117.18 | 178.44 | 58.83 | 38.19 |
| 18Val | None | None | None | None | None |
| 19Ile | None | None | None | None | None |
| 20Ile | None | None | None | None | None |
| 21Phe | None | None | None | None | None |
| 22Leu | None | None | 178.6 | 58.33 | 39.86 |
| 23Thr | 8.1 | 118.04 | 177.68 | 66.47 | 67.99 |
| 24Phe | 8.52 | 118.01 | 177.9 | 57.64 | 38.19 |
| 25Leu | None | None | None | None | None |
| 26Phe | None | None | None | None | None |
| 27Gln | None | None | None | None | None |
| 28His | None | None | None | None | None |
| 29Met | None | None | None | 58.32 | 31.77 |
| 30Phe | 8.29 | 116.33 | 175.92 | 59.69 | 38.76 |
| 31Ala | 8.76 | 121.72 | 177.75 | 55.58 | 17.4 |
| 32Ala | 8.09 | 120.7 | 178.89 | 54.83 | 17.49 |
| 33Thr | None | None | 179.25 | 66.68 | 68.49 |
| 34Leu | 8.76 | 123.36 | 178.94 | 57.34 | 39.68 |
| 35Lys | 8.1 | 117.81 | 178.51 | 58.92 | 31.38 |
| 36Glu | None | None | None | None | None |
| 37Gln | None | None | None | None | None |
| 38Ile | None | None | None | None | None |
| 39Gly | None | None | None | None | None |
| 40Met | None | None | None | None | None |
| 41Arg | None | None | None | None | None |
| 42Ala | 8.78 | 121.34 | 179.63 | 55.29 | 17.4 |
| 43Leu | 8.74 | 117.48 | 178.28 | 57.33 | 40.07 |
| 44Asn | 8.89 | 116.89 | 178.77 | 56.25 | 38.24 |
| 45Val | 8.15 | 121.01 | 177.3 | 67.41 | 30.75 |
| 46Ala | 8.84 | 123.99 | 179.76 | 55.6 | 17.33 |
| 47Glu | 9.13 | 114.35 | 179.94 | 58.23 | 29.28 |

| Residue | H | N | C | Ca | Cb |
|---|---|---|---|---|---|
| 48Thr | 8 | 117.58 | 177.66 | 66.88 | 68.17 |
| 49Val | 8.59 | 121.74 | 178.7 | 67.04 | 31.1 |
| 50Ala | 8.77 | 118.57 | 178.21 | 54.59 | 17.96 |
| 51Ser | 7.4 | 111.66 | 174.35 | 59.26 | 64.25 |
| 52Thr | 7.84 | 114.99 | 176 | 62.6 | 70.2 |
| 53Ser | None | None | None | None | None |
| 54Leu | None | None | 177.5 | 57.96 | 41.3 |
| 55Val | 6.91 | 114.61 | 175.88 | 65.07 | 30.26 |
| 56Arg | 6.84 | 115.07 | 180.15 | 59.19 | 29.11 |
| 57Glu | 8.54 | 118.2 | 179.77 | 58.67 | 28.57 |
| 58Ala | 8.41 | 121.48 | 179.41 | 53.81 | 17.26 |
| 59Phe | 7.55 | 114.31 | 177.08 | 61.21 | 38.22 |
| 60Arg | 7.8 | 116.85 | 176.82 | 55.93 | 29.47 |
| 61Asp | 7.44 | 120.92 | 176.11 | 53.74 | 40.5 |
| 62Ser | 8.69 | 116.04 | 175.53 | 61.01 | 61.95 |
| 63Asn | 8.12 | 114.06 | 174.17 | 49.56 | 37.99 |
| 64Pro | None | None | 176.1 | 64.4 | 32.68 |
| 65Ser | 7.55 | 107.32 | 175.97 | 61.89 | 63.67 |
| 66Val | 7.25 | 118.99 | 178.58 | 66.29 | 29.86 |
| 67Arg | 8.33 | 113.98 | 179.3 | 57.75 | 31.37 |
| 68Leu | 8 | 116.48 | 177.48 | 57.61 | 42.29 |
| 69Gln | 8.7 | 119.21 | 175.02 | 60.07 | 23.98 |
| 70Pro | None | None | 180.11 | 65.29 | 29.72 |
| 71Phe | 6.66 | 117 | 176.1 | 59.94 | 38.97 |
| 72Ala | 8.86 | 122.12 | 179.68 | 55.33 | 17.42 |
| 73Glu | 8.13 | 115.66 | 178.6 | 57.48 | 29.5 |
| 74Arg | 7.63 | 120.98 | 178.51 | 58.64 | 28.61 |
| 75Ile | 7.84 | 117.32 | 179.68 | 61.79 | 33.8 |
| 76Arg | 8.96 | 124.99 | 178.03 | 60.56 | 28.31 |
| 77Gln | 8.05 | 116.52 | 179.05 | 58.47 | 27.83 |
| 78Lys | None | None | None | None | None |
| 79Thr | None | None | 176.14 | 62.86 | 71.07 |
| 80Gly | 7.73 | 108.48 | 174.09 | 45.12 | None |
| 81Ala | 7.13 | 121.62 | 175.7 | 51.11 | 17.37 |
| 82Glu | 8.05 | 120.9 | 176.6 | 57.3 | 28.14 |
| 83Tyr | 6.41 | 104.33 | 172.77 | 54.45 | 38.38 |
| 84Val | 7.88 | 120.87 | 174.27 | 62.11 | 31.11 |
| 85Val | 9.79 | 126.78 | 175.96 | 60.58 | 33.4 |
| 86Ile | 7.49 | 124.28 | 175.4 | 59.42 | 39.37 |
| 87Gly | 9.78 | 115.73 | 172.9 | 43.77 | None |
| 88Asn | 7.99 | 116.41 | 176.54 | 50.67 | 38.28 |
| 89Arg | 7.42 | 115.49 | 176.19 | 58.2 | 29.03 |
| 90Gln | 7.35 | 115.41 | 176.08 | 55.49 | 27.85 |
| 91Gly | 8.22 | 107.74 | 172.72 | 45.12 | None |
| 92Ile | 7.5 | 119.84 | 175.83 | 57.89 | 35.23 |
| 93Ala | 7.77 | 125.43 | 177 | 52.13 | 18.11 |
| 94Tyr | 9.81 | 122.4 | 174.82 | 55.31 | 39.75 |
| 95Ala | 7.86 | 119.92 | 175.32 | 51.11 | 21.47 |
| 96His | 9.06 | 123.61 | 172.61 | 55.68 | 33.68 |

| Residue | H | N | C | Ca | Cb |
|---|---|---|---|---|---|
| 97Pro | None | None | 178.11 | 64.28 | 30.82 |
| 98Leu | 10.96 | 123.13 | 177.36 | 52.94 | 39.94 |
| 99Thr | 8.47 | 120.32 | 177.86 | 65.71 | 67.76 |
| 100Glu | 9.36 | 119.64 | 176.6 | 57.8 | 27.89 |
| 101Arg | 7.97 | 119.87 | 178.56 | 52.57 | 28.32 |
| 102Ile | 7.26 | 121.92 | 177.44 | 63.67 | 36.19 |
| 103Gly | 9.59 | 115.29 | 173.66 | 44.66 | None |
| 104Lys | 7.8 | 118.43 | 176.71 | 53.42 | 34.06 |
| 105Ser | 8.32 | 116.96 | 173.66 | 59.7 | 63.44 |
| 106Met | 8.75 | 122.42 | 176.49 | 56.29 | 34.91 |
| 107Ile | 10.24 | 129.59 | 175.34 | 59.45 | 39.08 |
| 108Gly | 8.27 | 112.21 | 176.19 | 44.67 | None |
| 109Gly | 8.95 | 108.53 | 174.98 | 46.28 | None |
| 110Asp | 8.13 | 116.43 | 176.62 | 52.02 | 37.97 |
| 111Asn | 8.38 | 115.1 | 177.77 | 53.95 | 38.46 |
| 112Lys | 7.9 | 120.01 | 179.01 | 60.81 | 31.65 |
| 113Glu | 9 | 115.4 | 178.97 | 58.23 | 28.47 |
| 114Val | 7 | 119.77 | 180.54 | 62.96 | 30.35 |
| 115Leu | 7.59 | 118.16 | 178.09 | 56.46 | 40.19 |
| 116Lys | 7.04 | 115.96 | 176.91 | 56.03 | 31.79 |
| 117Gly | 8.35 | 107.26 | 173.7 | 44.99 | None |
| 118Lys | 7.79 | 122.41 | 174.16 | 54.99 | 32.88 |
| 119Ser | 8.09 | 116.45 | 175.14 | 57.18 | 63.55 |
| 120Ile | 8.97 | 116.82 | 173.59 | 59.11 | 42.19 |
| 121Ile | 8.11 | 120.95 | 176.27 | 60.8 | 38.51 |
| 122Ser | 9.54 | 122.26 | 173.76 | 55.06 | 64.81 |
| 123Glu | 8.47 | 128.46 | 175.29 | 54.6 | 30.22 |
| 124Ala | 8.73 | 126.35 | 174.8 | 50.58 | 21.51 |
| 125Val | 8.56 | 119.8 | 175.53 | 61.92 | 29.94 |
| 126Gly | 6.75 | 112.11 | 176.93 | 44.29 | None |
| 127Ser | 7.22 | 120.8 | 176.75 | 62.72 | 61.94 |
| 128Leu | 11.08 | 123.33 | 176.8 | 54.75 | 40.71 |
| 129Gly | 7.2 | 104.79 | 172.21 | 43.64 | None |
| 130Pro | None | None | 177.26 | 63.36 | 31.3 |
| 131Ala | 9.49 | 129.45 | 175.12 | 52.02 | 24.05 |
| 132Ile | 8.54 | 117.7 | 174.07 | 59.48 | 39.96 |
| 133Arg | 9.4 | 122.87 | 175.28 | 53.99 | 33.01 |
| 134Gly | 9.15 | 110.62 | 171.98 | 43.73 | None |
| 135Lys | 9.17 | 122.8 | 173.66 | 54.07 | 33.22 |
| 136Ala | 8.91 | 122.74 | 173.91 | 47.51 | 21.16 |
| 137Pro | None | None | 174.64 | 60.64 | 30.5 |
| 138Ile | 7.52 | 118.12 | 175.38 | 59.52 | 38.78 |
| 139Phe | 8.75 | 126.04 | 177.13 | 56.42 | 41.53 |
| 140Asp | 8.93 | 119.03 | 178.3 | 51.65 | 40.59 |
| 141Glu | 9.58 | 117.95 | 176.78 | 58.59 | 27.97 |
| 142Asn | 8.21 | 116.08 | 175.86 | 52.49 | 39.02 |
| 143Gly | 8.04 | 107.8 | 174.35 | 45.48 | None |
| 144Ser | 8.63 | 118.73 | 174.08 | 57.61 | 62.98 |
| 145Val | 8.85 | 124.59 | 177.4 | 63.16 | 30.6 |

| Residue | H | N | C | Ca | Cb |
|---|---|---|---|---|---|
| 146Ile | 8.91 | 120.67 | 175.28 | 60.49 | 38.88 |
| 147Gly | 7.25 | 106.63 | 170.13 | 45.64 | None |
| 148Ile | 8.63 | 118.89 | 174.14 | 60.13 | 44.55 |
| 149Val | 9.14 | 124.78 | 174.31 | 60.67 | 32.75 |
| 150Ser | 9.55 | 122.29 | 173.9 | 54.98 | 65.04 |
| 151Val | 9.13 | 128.8 | 174.96 | 60.15 | 31.9 |
| 152Gly | 7.04 | 112.16 | 171.18 | 43.34 | None |
| 153Phe | 8.52 | 116.72 | 175.17 | 56.73 | 42.05 |
| 154Leu | 9.58 | 120.47 | 179.35 | 54.98 | 39.41 |
| 155Leu | 8.82 | 123.82 | 179.74 | 57.34 | 39.89 |
| 156Glu | 8.97 | 117.77 | 178.26 | 59.06 | 28.91 |
| 157Asp | 8.59 | 118.28 | 178.27 | 57 | 40.77 |
| 158Ile | 8.6 | 121.66 | 177.72 | 65.97 | 37.7 |
| 159Gln | None | None | None | None | None |
| 160Arg | None | None | 177.45 | 60.36 | 28.92 |
| 161Thr | 8.4 | 120.11 | 177.77 | 65.52 | 67.75 |
| 162Val | 7.65 | 118.22 | 177.36 | 66.92 | 30.89 |
| 163Trp | None | None | None | None | None |
| 164Ser | None | None | None | None | None |
| 165Tyr | None | None | None | None | None |
| 166Ser | None | None | None | None | None |
| 167Met | None | None | None | None | None |
| 168Lys | None | None | None | None | None |
| 169Ile | None | None | None | None | None |
| 170Phe | None | None | None | None | None |
| 171Phe | None | None | None | None | None |
| 172Phe | None | 122.24 | 176.11 | 58.93 | 38.84 |
| 173Ser | 8.85 | 121.35 | 178.72 | 59.09 | 61.72 |
| 174Val | 8.39 | 120.86 | 177.52 | 65.82 | 28.35 |
| 175Leu | 8.66 | 120.44 | 177.54 | 60.18 | 41.06 |
| 176Ala | 8.43 | 119.78 | 178.31 | 55.24 | 17.21 |
| 177Leu | 8.5 | 117.59 | 177.93 | 57.7 | 40.3 |
| 178Leu | 8.33 | 119.16 | 178.47 | 57.8 | 40.13 |
| 179Phe | 8.28 | 116 | 176.72 | 57.72 | 38.22 |
| 180Gly | 8.36 | 107.77 | 174.7 | 45.21 | None |
| 181Ala | 8.67 | 118.85 | 175.03 | 54.81 | 18.02 |
| 182Val | 9.58 | 122.23 | 173.93 | 64.64 | 32.86 |
| 183Gly | 7.63 | 107.63 | 172.76 | 45.31 | None |
| 184Ala | 8.19 | 121.62 | 177.5 | 54.83 | 17.37 |
| 185Val | 8.01 | 121.1 | 177.37 | 67.44 | 30.86 |
| 186Ala | 8.83 | 124.02 | 177.95 | 55.38 | 17.28 |
| 187Ile | 7.86 | 116.29 | 177.66 | 66.28 | 38.54 |
| 188Ala | 8.77 | 121.79 | 179.6 | 55.39 | 17.55 |
| 189Lys | 8.34 | 114.84 | 179.01 | 57.53 | 29.92 |
| 190Ala | 7.67 | 121.33 | 177.55 | 55.04 | 17.68 |
| 191Val | 9.03 | 115.37 | 177.76 | 58.28 | 28.25 |
| 192Lys | 8.02 | 115.87 | 176.04 | 58.23 | 30.73 |
| 193Lys | 7.77 | 118.84 | 178.6 | 54.77 | 34.16 |
| 194Ser | None | None | None | None | None |

| Residue | H | N | C | Ca | Cb |
|---|---|---|---|---|---|
| 195Ile | None | None | None | None | None |
| 196His | None | None | None | None | None |
| 197Gly | None | None | None | None | None |
| 198Leu | None | None | None | None | None |
| 199Glu | None | None | None | None | None |
| 200Pro | None | None | 178.45 | 66.72 | 29.05 |
| 201Glu | 8.5 | 117.56 | 178.11 | 57.49 | 28.47 |
| 202Glu | 7.98 | 121.27 | 177.42 | 57.39 | 29.26 |
| 203Ile | 7.65 | 121.01 | 177.72 | 64.63 | 41.29 |
| 204Gly | 8.37 | 107.02 | 174.86 | 47.38 | None |
| 205Leu | 8.36 | 119.26 | 177.72 | 57.81 | 40.17 |
| 206Leu | None | None | None | None | None |
| 207Tyr | None | None | 177.66 | 58.38 | 38.79 |
| 208Gln | 7.89 | 117.89 | 179.01 | 57.47 | 29.59 |
| 209Glu | 7.75 | 120.82 | 178.55 | 57.4 | 28.56 |
| 210Lys | 7.94 | 117.23 | 177.75 | 58.46 | 31.34 |
| 211Gln | 7.84 | 116.63 | 178.96 | 58.65 | 28.08 |
| 212Ala | None | None | None | None | None |
| 213Ile | None | None | 179.19 | None | None |
| 214Leu | 8.55 | 122.95 | None | 57.64 | 40.31 |
| 215Glu | 7.74 | 114.79 | 178.34 | 57.48 | 28.46 |
| 216Ala | 8.08 | 121.32 | 179.49 | 55.12 | 17.56 |
| 217Ile | 8.12 | 118.39 | 178.65 | 66.54 | 36.33 |
| 218Arg | None | None | None | None | None |
| 219Glu | None | None | None | None | None |
| 220Gly | None | None | None | None | None |
| 221Ile | None | None | None | None | None |
| 222Val | None | None | None | None | None |
| 223Ala | None | None | None | None | None |
| 224Ile | None | None | None | None | None |
| 225Asn | None | None | None | None | None |
| 226Gln | None | None | None | None | None |
| 227Glu | None | None | None | None | None |
| 228Gly | None | None | None | None | None |
| 229Thr | 7.53 | 114.85 | 174.53 | 61.07 | 68.18 |
| 230Ile | 8.33 | 124.24 | 177.27 | 62.71 | 36.12 |
| 232Met | None | None | None | None | None |
| 233Val | None | None | None | None | None |
| 234Asn | None | None | None | None | None |
| 235Gln | None | None | None | None | None |
| 236Thr | None | None | 178.53 | 65.85 | 68.4 |
| 237Ala | 8.3 | 120.01 | 179.67 | 55.18 | 17.16 |
| 238Leu | 8.61 | 117.5 | 179.51 | 57.47 | 40.06 |
| 239Lys | None | None | None | None | None |
| 240Leu | None | None | None | None | None |
| 241Leu | None | None | None | None | None |
| 242Gly | None | None | None | None | None |
| 243Tyr | None | None | None | None | None |
| 244Asp | None | None | None | None | None |

| Residue | H | N | C | Ca | Cb |
|---|---|---|---|---|---|
| 245Asn | None | None | None | None | None |
| 246Glu | None | None | None | None | None |
| 247Arg | None | None | None | None | None |
| 248Asn | None | None | None | None | None |
| 249Val | None | None | None | None | None |
| 250Leu | None | None | 178.08 | None | 40.4 |
| 251Gly | 8.95 | 109.6 | 174.17 | 44.86 | None |
| 252Thr | None | None | None | None | None |
| 253Pro | None | None | None | None | None |
| 254Ile | None | None | None | None | None |
| 255Leu | None | None | None | None | None |
| 256Gln | None | None | None | None | None |
| 257Leu | None | None | None | None | None |
| 258Ile | None | None | None | None | None |
| 259Pro | None | None | None | None | None |
| 260His | None | None | None | None | None |
| 261Ser | None | None | None | None | None |
| 262Arg | None | None | None | None | None |
| 263Leu | None | None | None | None | None |
| 264Pro | None | None | None | None | None |
| 265Glu | None | None | None | None | None |
| 266Val | None | None | None | None | None |
| 267Ile | None | None | None | None | None |
| 268Arg | None | None | None | None | None |
| 269Thr | None | None | None | None | None |
| 270Gly | None | None | 172.83 | 47.23 | None |
| 271Gln | 8.03 | 119.23 | 174.35 | 53.62 | 29.3 |
| 272Ala | 8.38 | 125.14 | 176.44 | 51.72 | 20 |
| 273Glu | 8.56 | 118.25 | 177.78 | 57.5 | 28.78 |
| 274Tyr | 7.64 | 121.06 | 177.48 | 57.74 | 41.29 |
| 275Asp | 8.95 | 119.06 | 175.43 | 51.66 | 40.67 |
| 276Asp | None | None | None | None | None |
| 277Glu | None | None | None | None | None |
| 278Met | 9.13 | 124.81 | 174.97 | 54.49 | 30.62 |
| 279Val | None | None | None | None | None |
| 280Leu | None | None | None | None | None |
| 281Gly | None | None | None | None | None |
| 282Gly | None | 104.25 | 173.66 | 44.73 | None |
| 283Glu | 7.79 | 121.01 | 174.31 | 54.81 | 30.9 |
| 284Thr | 8.66 | 118.52 | 175.93 | 62.6 | 70.69 |
| 285Val | 9.14 | 121.46 | 177.87 | 58.87 | 35.63 |
| 286Ile | None | None | None | None | None |
| 287Ala | None | None | None | None | None |
| 288Asn | None | None | None | None | None |
| 289Arg | None | None | None | None | None |
| 290Ile | None | None | None | None | None |
| 291Pro | None | None | None | None | None |
| 292Ile | None | None | None | None | None |
| 293Lys | 8.57 | 126.26 | 176.02 | 54.06 | 35.29 |

| Residue | H | N | C | Ca | Cb |
|---|---|---|---|---|---|
| 294Asn | 8.46 | 117.91 | None | 50.23 | None |
| 295Lys | None | None | 177.28 | 58.32 | 38.24 |
| 296Gln | 7.43 | 115.93 | 176.25 | 55.63 | 28.94 |
| 297Gly | 8.2 | 107.55 | 173.8 | 45.23 | None |
| 298Arg | 7.74 | 121.97 | 175.95 | 54.84 | 32.79 |
| 299Val | None | None | None | None | None |
| 300Ile | None | None | None | None | None |
| 301Gly | None | None | 171.34 | 45.25 | None |
| 302Ala | 8.51 | 120.02 | 174.2 | 52.15 | 21.87 |
| 303Val | None | None | None | None | None |
| 304Ser | None | None | None | None | None |
| 305Thr | None | None | None | None | None |
| 306Phe | None | None | None | None | None |
| 307Arg | None | None | None | None | None |
| 308Asn | None | None | None | None | None |
| 309Lys | None | None | None | None | None |

**Table S2. Chemical shifts from $^{1}H$ -detected sequence specific assignment of the citrate free state of CitApc.**

| Residue | H | N | C | Ca | Cb |
|---|---|---|---|---|---|
| 1Met | None | None | None | None | None |
| 2Lys | None | None | None | None | None |
| 3Leu | None | None | None | None | None |
| 4Gln | None | None | None | None | None |
| 5Thr | None | None | 174.49 | None | None |
| 6Arg | 9.33 | 119.99 | 176.4 | 54.25 | 30.7 |
| 7Leu | 8.78 | 120.72 | 177.76 | 58.95 | 40.89 |
| 8Met | 8.36 | 119.98 | 177.83 | 58.14 | 30.85 |
| 9Val | 7.81 | 118.1 | 177.63 | 66.51 | 30.73 |
| 10Ile | 8.85 | 121.7 | 177.52 | 66.97 | 38.88 |
| 11Ile | 8.83 | 122.5 | 178.09 | 66.95 | 36.28 |
| 12Ala | 8.65 | 122.23 | 178.09 | 55.32 | 17.38 |
| 13Ser | 8.58 | 118.39 | 178.25 | 59.76 | 62.85 |
| 14Leu | 8.45 | 119.21 | 178.02 | 57.78 | 40.87 |
| 15Leu | 7.98 | 120.37 | 178.99 | 55.27 | 40.75 |
| 16Leu | 8.25 | 118.28 | 178.44 | 58.46 | 38.33 |
| 17Phe | 8.29 | 118.36 | None | None | None |
| 18Val | None | None | None | None | None |
| 19Ile | None | None | None | None | None |
| 20Ile | None | None | 175.12 | 63.04 | 41.42 |
| 21Phe | 8.78 | 120.32 | 179.17 | 58.72 | 38.37 |
| 22Leu | 8.01 | 117.1 | 179.03 | 57.7 | 40.29 |
| 23Thr | 8.05 | 118.3 | 179.11 | 66.88 | 68.11 |
| 24Phe | 8.57 | 122.24 | 177.87 | 60.14 | 38.54 |
| 25Leu | 7.98 | 117.65 | 178.48 | 57.23 | 38.39 |
| 26Phe | 8.78 | 121.76 | 178.28 | 59.81 | 37.57 |
| 27Gln | 8.47 | 118.99 | 178.98 | 58.95 | 28.39 |
| 28His | 8.25 | 118.19 | 178.65 | 59.67 | 30.23 |
| 29Met | 7.53 | 117.92 | 178.92 | 58.98 | 30.27 |
| 30Phe | 7.89 | 118.47 | 177.88 | 60.44 | 40.15 |
| 31Ala | 8.58 | 120.5 | 178.07 | 54.91 | 16.58 |
| 32Ala | 8.54 | 120.06 | 178.51 | 55.39 | 17.14 |
| 33Thr | 7.94 | 115.01 | 178.89 | 57.5 | 68.36 |
| 34Leu | 7.75 | 120.78 | 178.45 | 57.78 | 40.67 |
| 35Lys | None | None | None | None | None |
| 36Glu | None | None | None | None | None |
| 37Gln | None | None | None | None | None |
| 38Ile | None | None | None | None | None |
| 39Gly | None | None | None | None | None |
| 40Met | None | None | None | None | None |
| 41Arg | None | None | None | None | None |
| 42Ala | None | None | None | None | None |
| 43Leu | None | None | None | None | None |
| 44Asn | None | None | None | None | None |
| 45Val | None | None | None | None | None |
| 46Ala | None | None | 179.29 | 55.92 | None |

| Residue | H | N | C | Ca | Cb |
|---|---|---|---|---|---|
| 47Glu | 9.19 | 114.54 | 179.11 | 58.36 | 28.83 |
| 48Thr | None | None | None | 66.85 | 68.19 |
| 49Val | 8.66 | 122.01 | 178.12 | 67.02 | 30.77 |
| 50Ala | 8.58 | 118.83 | 178.06 | 54.67 | 18.44 |
| 51Ser | 7.44 | 109.79 | 178.31 | 58.15 | 63.75 |
| 52Thr | None | None | None | None | None |
| 53Ser | None | None | None | 62.51 | None |
| 54Leu | 7.88 | 120.17 | 178.05 | 58.1 | 40.78 |
| 55Val | 6.97 | 116.5 | 176.44 | 65.54 | 30.24 |
| 56Arg | 7.03 | 116 | 179.93 | 59.46 | 29.13 |
| 57Glu | 8.28 | 118.07 | 179.11 | 58.59 | 30.77 |
| 58Ala | 8.51 | 122.13 | 179.62 | 53.97 | 17.22 |
| 59Phe | 7.64 | 114.71 | 177.15 | 61.23 | 38.42 |
| 60Arg | 7.82 | 116.83 | 176.81 | 55.87 | 29.49 |
| 61Asp | 7.46 | 121.1 | 176.09 | 53.87 | 40.51 |
| 62Ser | 8.73 | 116.05 | 175.57 | 61.1 | 61.93 |
| 63Asn | 8.19 | 114.12 | 174.18 | 49.58 | 37.93 |
| 64Pro | None | None | 176.1 | 64.27 | None |
| 65Ser | 7.6 | 107.49 | 175.97 | 61.8 | 63.56 |
| 66Val | 7.26 | 119.68 | 178.56 | 66.26 | 29.93 |
| 67Arg | 8.27 | 113.92 | 179.55 | 57.65 | 31.52 |
| 68Leu | 8.07 | 116.56 | 177.42 | 57.36 | 41.99 |
| 69Gln | 8.73 | 119.25 | 174.19 | 54.78 | 30.94 |
| 70Pro | None | 118.74 | 179.88 | 65.06 | 29.7 |
| 71Phe | 6.7 | 117.81 | 176.17 | 60.31 | 39.01 |
| 72Ala | 8.87 | 121.71 | 179.45 | 55.36 | 17.45 |
| 73Glu | 8.13 | 115.97 | 177.65 | 58.03 | 27.96 |
| 74Arg | 8.01 | 120.11 | 177.99 | 58.03 | 29.24 |
| 75Ile | 8.07 | 118.8 | 179.66 | 62.42 | 40.82 |
| 76Arg | 8.85 | 124.61 | 178.1 | 60.42 | 30.37 |
| 77Gln | 8.04 | 116.73 | 178.9 | 58.43 | 27.81 |
| 78Lys | 8.14 | 117.28 | 178.38 | 57.39 | 32.23 |
| 79Thr | 7.89 | 104.15 | 176.98 | 62.07 | 70.07 |
| 80Gly | 8.56 | 109.72 | 174.13 | 45.59 | None |
| 81Ala | 7.32 | 122.12 | 176.16 | 51.46 | 17.7 |
| 82Glu | 8.08 | 120.81 | 176.2 | 57.51 | 29.21 |
| 83Tyr | 6.46 | 104.25 | 172.88 | 54.39 | 38.61 |
| 84Val | 7.87 | 120.25 | 173.92 | 62.04 | 32.49 |
| 85Val | 9.5 | 126.7 | 175.41 | 59.88 | 33.9 |
| 86Ile | 8.08 | 123.98 | 175.44 | 59.77 | 40.22 |
| 87Gly | 9.79 | 115.65 | 172.69 | 43.89 | None |
| 88Asn | 7.99 | 116.52 | 176.66 | 50.8 | 38.4 |
| 89Arg | 7.48 | 115.73 | 176.08 | 58.19 | 28.87 |
| 90Gln | 7.48 | 115.76 | 176.06 | 55.97 | 28.17 |
| 91Gly | 8.2 | 107.17 | 172.76 | 45.16 | None |
| 92Ile | 7.5 | 119.79 | 175.96 | 57.88 | 35.3 |
| 93Ala | 7.83 | 125.53 | 177.04 | 52.3 | 18.12 |
| 94Tyr | 9.82 | 122.45 | 174.5 | 55.18 | 39.78 |
| 95Ala | 7.92 | 119.54 | 175.46 | 51.03 | 21.53 |

| Residue | H | N | C | Ca | Cb |
|---|---|---|---|---|---|
| 96His | 9.24 | 124.54 | 173.7 | 54.25 | 33.08 |
| 97Pro | None | None | None | None | None |
| 98Leu | 10.98 | 123.02 | 177.36 | 52.94 | None |
| 99Thr | None | None | None | None | None |
| 100Glu | None | None | None | 58.5 | 27.96 |
| 101Arg | 8.33 | 116.44 | 179.03 | 55.61 | 28.64 |
| 102Ile | 7.31 | 121.6 | 177.67 | 63.71 | 36.14 |
| 103Gly | 9.56 | 115.5 | 172.46 | 44.75 | None |
| 104Lys | 7.88 | 118.61 | 174.27 | 53.7 | 33.72 |
| 105Ser | None | None | None | None | None |
| 106Met | None | None | None | None | None |
| 107Ile | None | None | None | None | None |
| 108Gly | 9.63 | None | 173.48 | 44.85 | None |
| 109Gly | 9.03 | 108.75 | 172.98 | 46.34 | None |
| 110Asp | 8 | 118.91 | 176.91 | 53.73 | None |
| 111Asn | None | None | None | None | None |
| 112Lys | None | None | None | None | None |
| 113Glu | None | None | 178.97 | 58.69 | None |
| 114Val | 7.04 | 119.65 | 177.94 | 63.07 | 30.59 |
| 115Leu | 7.69 | 118.15 | 178.06 | 56.48 | 40.27 |
| 116Lys | 7.1 | 116.3 | 176.93 | 56.42 | 31.84 |
| 117Gly | 8.43 | 106.98 | 173.51 | 45.08 | None |
| 118Lys | 7.73 | 121.93 | 174.26 | 55.14 | 29.31 |
| 119Ser | 8.22 | 116.43 | 178.08 | 57.54 | None |
| 120Ile | None | None | None | None | None |
| 121Ile | None | None | None | None | None |
| 122Ser | None | None | None | None | None |
| 123Glu | None | None | 173.07 | 59.68 | 31.58 |
| 124Ala | 8.81 | 126.28 | 177.19 | 50.53 | 19.42 |
| 125Val | 8.85 | 119.33 | 178.22 | 60.25 | 31.79 |
| 126Gly | 7.24 | 112.29 | 171.5 | 43.36 | None |
| 127Ser | None | None | None | None | None |
| 128Leu | None | None | None | None | None |
| 129Gly | None | None | None | None | None |
| 130Pro | None | None | None | None | None |
| 131Ala | None | None | None | None | None |
| 132Ile | None | None | 175.35 | 60.31 | None |
| 133Arg | 9.14 | 125.86 | 174.72 | 53.88 | 32.54 |
| 134Gly | 9.18 | 109.99 | 171.69 | 43.67 | None |
| 135Lys | 9.07 | 122.9 | 173.85 | 54.17 | 36.43 |
| 136Ala | 9.01 | 122.27 | 173.84 | 47.79 | 21.88 |
| 137Pro | None | None | 174.41 | 60.61 | 30.9 |
| 138Ile | 7.52 | 117.86 | 175.16 | 59.66 | 38.81 |
| 139Phe | 8.88 | 126.13 | 177.14 | 56.5 | 41.55 |
| 140Asp | 8.96 | 119.13 | 178.35 | 51.7 | 40.6 |
| 141Glu | 9.56 | 117.86 | 176.71 | 58.6 | 27.94 |
| 142Asn | 8.23 | 116.07 | 175.83 | 52.61 | 38.98 |
| 143Gly | 8.07 | 107.87 | 174.33 | 45.5 | None |
| 144Ser | 8.64 | 118.67 | 173.95 | 57.71 | 63 |

| Residue | H | N | C | Ca | Cb |
|---|---|---|---|---|---|
| 145Val | 8.91 | 124.54 | 177.55 | 63.17 | 30.54 |
| 146Ile | 8.95 | 120.57 | 175.47 | 60.59 | 38.84 |
| 147Gly | 7.28 | 106.82 | 169.96 | 45.69 | None |
| 148Ile | 8.59 | 118.29 | 174.08 | 60.07 | 44.54 |
| 149Val | 9.15 | 124.88 | 174.36 | 60.54 | 33.08 |
| 150Ser | 9.41 | 121.27 | 173.68 | 55.18 | 64.54 |
| 151Val | 9.18 | 126.91 | 174.24 | 60.21 | 33.28 |
| 152Gly | 7.14 | 112.07 | 171.8 | 43.41 | None |
| 153Phe | 8.41 | 117.96 | 178.54 | 55.98 | 40.66 |
| 154Leu | 8.92 | 120.34 | 174.83 | 54.27 | 40.98 |
| 155Leu | 8.91 | 120.64 | 174.87 | 55.22 | 40.97 |
| 156Glu | 8.64 | 120.42 | 176.56 | 54.24 | 30.94 |
| 157Asp | 7.91 | 120.42 | 175.54 | 55.07 | 40.74 |
| 158Ile | 8.63 | 119.09 | 175.29 | 60.01 | 37.29 |
| 159Gln | None | None | None | None | None |
| 160Arg | None | None | None | None | None |
| 161Thr | None | None | None | None | None |
| 162Val | None | None | None | None | None |
| 163Trp | None | None | None | None | None |
| 164Ser | None | None | None | None | None |
| 165Tyr | None | None | None | None | None |
| 166Ser | None | None | None | None | None |
| 167Met | None | None | None | None | None |
| 168Lys | None | None | None | None | None |
| 169Ile | None | None | None | None | None |
| 170Phe | None | None | None | None | None |
| 171Phe | None | None | None | None | None |
| 172Phe | None | None | None | 60.28 | 37.69 |
| 173Ser | 8.75 | 121.59 | 174.43 | 60.06 | None |
| 174Val | 8.77 | 121.26 | 177.76 | 66.81 | 30.84 |
| 175Leu | 8.69 | 119.73 | 178.21 | 60.09 | 40.99 |
| 176Ala | None | None | None | None | None |
| 177Leu | None | None | None | None | None |
| 178Leu | None | None | None | None | None |
| 179Phe | None | None | 175.13 | None | None |
| 180Gly | 8.27 | 112.13 | 176.29 | 44.65 | None |
| 181Ala | 8.18 | 116.72 | 179.34 | 52.4 | 21.6 |
| 182Val | 9.08 | 124.39 | 174.61 | 60.46 | 33.14 |
| 183Gly | 9.14 | 110.32 | 171.48 | 43.67 | None |
| 184Ala | 8.87 | 122.78 | 179.34 | 54.95 | 17.18 |
| 185Val | 8.1 | 116.76 | 177.42 | 66.41 | 28.76 |
| 186Ala | 8.71 | 122.44 | 179.8 | 54.67 | 17.23 |
| 187Ile | 8.24 | 117.96 | 177.4 | 66.24 | 38.36 |
| 188Ala | 8.84 | 122.2 | 179.42 | 55.85 | 17.29 |
| 189Lys | 9.12 | 115.65 | 180.12 | 59.13 | 29.02 |
| 190Ala | 8.59 | 118.57 | 179.29 | 55.32 | 17.82 |
| 191Val | 7.77 | 116.57 | 177.46 | 66.17 | 31.08 |
| 192Lys | 8.3 | 119.45 | 178.15 | 58.84 | 30.93 |
| 193Lys | 8.44 | 121.15 | 178.88 | 59.97 | 30.57 |

| Residue | H | N | C | Ca | Cb |
|---|---|---|---|---|---|
| 194Ser | None | None | 174.29 | 61.63 | 68.38 |
| 195Ile | 8.32 | 124.4 | 176.88 | 62.53 | 37.66 |
| 196His | 8.64 | 117.41 | 177.69 | 62.33 | 28.26 |
| 197Gly | 8.52 | 107.35 | 175.15 | 45.05 | None |
| 198Leu | 7.96 | 123.81 | 175.39 | 59.75 | 40.24 |
| 199Glu | None | None | None | None | None |
| 200Pro | None | None | 177.75 | 66.62 | 30.85 |
| 201Glu | 8.58 | 118.95 | 178.95 | 57.71 | 30.97 |
| 202Glu | 7.84 | 120.65 | 177.71 | 57.81 | 29.45 |
| 203Ile | 7.76 | 120.62 | 179.28 | 62.06 | 40.82 |
| 204Gly | 8.63 | 107.48 | 176.31 | 45.03 | None |
| 205Leu | 7.66 | 121.78 | 178.61 | 55.18 | 40.61 |
| 206Leu | 8.07 | 116.7 | 177.52 | 57.44 | 38.24 |
| 207Tyr | 8.98 | 119.48 | 178.58 | 59 | 38.31 |
| 208Gln | 8.07 | 117.51 | 177.61 | 57.76 | 27.99 |
| 209Glu | 7.99 | 119.96 | 178.7 | 59.48 | 30.67 |
| 210Lys | 8.5 | 117.82 | 178.45 | 59.24 | 30.78 |
| 211Gln | 7.55 | 117.92 | 178.31 | 58.55 | 28.39 |
| 212Ala | 8.3 | 120.3 | 179.51 | 55.24 | 17.41 |
| 213Ile | 8.27 | 118.48 | 177.52 | 66.59 | 38.27 |
| 214Leu | 8.31 | 118.67 | 177.96 | 57.53 | 41.04 |
| 215Glu | 8.82 | 120.21 | 176.3 | 57.18 | 29.5 |
| 216Ala | 7.72 | 120.11 | 178.14 | 54.42 | 17.34 |
| 217Ile | 8.18 | 117.62 | 177.64 | 66.73 | 38.47 |
| 218Arg | 8.35 | 121.51 | 175.76 | 59.91 | 30.64 |
| 219Glu | 8.46 | 119.36 | 179.11 | 55.43 | 29.09 |
| 220Gly | None | None | None | None | None |
| 221Ile | None | None | None | None | None |
| 222Val | 8.62 | 121.58 | 173.86 | 59.83 | 31.29 |
| 223Ala | 8.95 | 127.37 | 176.55 | 49.74 | 22.67 |
| 224Ile | 8.87 | 111.03 | 176.82 | 58 | 41.78 |
| 225Asn | 8 | 119.48 | 179.6 | 55.19 | 41.01 |
| 226Gln | None | None | 176.69 | 58.47 | 27.82 |
| 227Glu | 7.56 | 116.39 | 177.07 | 55.9 | 28.95 |
| 228Gly | 8.44 | 107.02 | 173.58 | 45.1 | None |
| 229Thr | 7.92 | 118.47 | 174.63 | 61.57 | 69.48 |
| 230Ile | 8.32 | 124.32 | 176.77 | 62.56 | 37.26 |
| 231Thr | 7.82 | 121.32 | None | None | None |
| 232Met | None | None | None | None | None |
| 233Val | None | None | None | None | None |
| 234Asn | None | None | None | None | None |
| 235Gln | 8.8 | 119.57 | 177.22 | 58.41 | 30.93 |
| 236Thr | 7.87 | 115.37 | 176.67 | 66.4 | 68.04 |
| 237Ala | 7.86 | 121.8 | 176.03 | 55.23 | 17.43 |
| 238Leu | 8.48 | 117.44 | 177.95 | 57.59 | 40.55 |
| 239Lys | 8.59 | 118.2 | 179.68 | 59.19 | 30.96 |
| 240Leu | 8.35 | 118.81 | 178.52 | 55.43 | 40.16 |
| 241Leu | 7.9 | 116.4 | 176.88 | 56.13 | 40.61 |
| 242Gly | 8.36 | 106.73 | 172.75 | 45.08 | None |

| Residue | H | N | C | Ca | Cb |
|---|---|---|---|---|---|
| 243Tyr | 7.6 | 120.37 | 176.58 | 58.09 | 40.66 |
| 244Asp | 8.47 | 117.89 | 178.03 | 54.79 | 41.73 |
| 245Asn | 7.65 | 115.66 | 178.9 | 55.74 | 38.72 |
| 246Glu | None | None | None | None | None |
| 247Arg | None | None | None | None | None |
| 248Asn | None | None | None | None | None |
| 249Val | None | None | None | None | None |
| 250Leu | 7.48 | 120.7 | 178.03 | 57.08 | 40.67 |
| 251Gly | 8.92 | 109.6 | 173.81 | 44.85 | None |
| 252Thr | 7.83 | 118.64 | 176.41 | 62.24 | None |
| 253Pro | None | 124.58 | 179.54 | 62.59 | 31.81 |
| 254Ile | 8.99 | 127.21 | 175.92 | 63.63 | 34.17 |
| 255Leu | 8.42 | 119.28 | 178.53 | 54.64 | 39.75 |
| 256Gln | 7.57 | 114.76 | 176.44 | 57.43 | 28.38 |
| 257Leu | 7.91 | 117.72 | 177.57 | 57.99 | 40.05 |
| 258Ile | 8.39 | 117.27 | 179.16 | 57.66 | 38.17 |
| 259Pro | None | None | 177.65 | 66.39 | 30.29 |
| 260His | 8.05 | 118.78 | 177.61 | 55.31 | 27.83 |
| 261Ser | None | None | None | 63.19 | None |
| 262Arg | 7.76 | 120.63 | 178.57 | 58.53 | 28.84 |
| 263Leu | 8.01 | 117.76 | 177.68 | 59.68 | 39.89 |
| 264Pro | None | None | 178.83 | 65.49 | 30.2 |
| 265Glu | 7.3 | 117.72 | 178.36 | 58.71 | 29.78 |
| 266Val | 8.32 | 119.25 | 177.31 | 65.39 | 29.59 |
| 267Ile | 8.73 | 117.69 | 176.99 | 62.7 | 38.8 |
| 268Arg | 7.28 | 117.56 | 177.16 | 56.54 | 30.04 |
| 269Thr | 8.56 | 107.04 | 177.27 | 62.21 | 71.54 |
| 270Gly | 8.42 | 111.5 | 173.01 | 46.35 | None |
| 271Gln | 8.04 | 119.15 | 174.36 | 53.62 | 29.6 |
| 272Ala | 8.47 | 125.01 | 176.79 | 51.7 | 20.16 |
| 273Glu | 8.78 | 119.34 | 174.62 | 59.6 | 30.76 |
| 274Tyr | 8.57 | 119.76 | 177.52 | 56.16 | 40.85 |
| 275Asp | 8.86 | 118.85 | 174.95 | 54.79 | 40.62 |
| 276Asp | 8.06 | 120.54 | None | 54.88 | None |
| 277Glu | None | None | None | None | None |
| 278Met | None | None | None | None | None |
| 279Val | None | None | None | None | None |
| 280Leu | None | None | 178.66 | 55.48 | None |
| 281Gly | 8.25 | 112.22 | 176.25 | 44.67 | None |
| 282Gly | 7.73 | 107.65 | 170.82 | 45.36 | None |
| 283Glu | 7.9 | 121.05 | 175.21 | 55.1 | 30.8 |
| 284Thr | 8.01 | 118.96 | 176.85 | 61.46 | 68.82 |
| 285Val | 7.47 | 121.37 | 176.07 | 58.95 | 35.65 |
| 286Ile | 7.79 | 117.79 | 175.59 | 59.42 | 40.25 |
| 287Ala | 9.26 | 130.53 | 176.6 | 49.97 | 21.73 |
| 288Asn | None | None | None | None | None |
| 289Arg | None | None | None | None | None |
| 290Ile | None | None | None | None | None |
| 291Pro | None | None | 174.9 | 62.78 | 30.69 |

| Residue | H | N | C | Ca | Cb |
|---|---|---|---|---|---|
| 292Ile | 8.35 | 121.4 | 175.69 | 59.24 | 38.69 |
| 293Lys | 8.83 | 125.52 | 175.95 | 53.87 | 35.8 |
| 294Asn | 8.56 | 117.93 | 178.01 | 50.27 | 37.88 |
| 295Lys | 8.32 | 117.53 | 177.77 | 58.5 | 27.88 |
| 296Gln | 7.55 | 116.11 | 176.13 | 55.87 | 28.87 |
| 297Gly | 8.17 | 107.52 | 173.55 | 45.19 | None |
| 298Arg | 8.14 | 120.94 | 176.27 | 54.89 | 30.23 |
| 299Val | 8.86 | 124.73 | 176.62 | 62.95 | 30.34 |
| 300Ile | 8.86 | 120.02 | 175.55 | 60.18 | 38.52 |
| 301Gly | 7.94 | 107.43 | 170.67 | 45.37 | None |
| 302Ala | 8.56 | 119.68 | 173.93 | 51.76 | 21.81 |
| 303Val | 8.92 | 118.9 | None | 58.83 | None |
| 304Ser | None | None | None | None | None |
| 305Thr | None | None | None | None | None |
| 306Phe | None | None | None | None | None |
| 307Arg | None | None | None | None | None |
| 308Asn | None | None | None | None | None |
| 309Lys | None | None | None | None | None |

**Table S3. NMR experimental parameters used in sequence specific assignment in the citrate free state of CitApc.** The powers for soft pulses are taken at the maximum value.

| Spectrum | (H)NH | (H)CANH | (H)(CO)CA(CO)NH | (H)CONH | (H)CO(CA)NH |
|---|---|---|---|---|---|
| Scans per point | 8 | 24 | 96 | 8 | 96 |
| Experimental time / h | 0.9 | 112 | 205.3 | 17.2 | 206.4 |
| Signal/noise | 85.99 | 28.22 | 10.29 | 20.85 | 15.06 |
| Field | 850 | 850 | 850 | 850 | 850 |
| Spinning frequency/Hz | 55000 | 55000 | 55000 | 55000 | 55000 |
| interscan delay /s | 1 | 1 | 1 | 1 | 1 |
| Sweep width (t1) / ppm | 160 (N) | 32 (N) | 40 (N) | 40 (N) | 40 (N) |
| Max indirect evolution (t1) / ms | 14.5 (N) | 21.8 (N) | 10.1 (N) | 12.5 (N) | 12.5 (N) |
| Sweep width (t2) / ppm | 40.3 (H) | 32 (C) | 30 (C) | 18.0 (C) | 18.0 (C) |
| Max indirect evolution (t2) / ms | 30.0 (H) | 10.2 (C) | 8.6 (C) | 11.7 (C) | 11.7 (C) |
| Sweep width (t3) / ppm | N/A | 58.8 (H) | 40.3 (H) | 40.3 (H) | 40.3 (H) |
| Max indirect evolution (t3) / ms | N/A | 20.5 (H) | 30.0 (H) | 30.0 (H) | 30.0 (H) |
| **Transfer I** | **HN (dipolar)** | **HCA (dipolar)** | **HCO (dipolar)** | **HCO (dipolar)** | **HCO (dipolar)** |
| $^{1}$H field / kHz | 99.4 | 95.1 | 96.3 | 100 | 100 |
| X field / kHz | 38.1 | 40.4 | 40.1 | 40.1 | 40.1 |
| Shape | 100 - 80 % ramp (H) | 85 - 100 % ramp (H) | 85 - 100 % ramp (H) | 85 - 100 % ramp (H) | 85 - 100 % ramp (H) |
| Carrier $^{13}$C | N/A | 53.7 | 173.3 | 173.8 | 173.8 |
| Time / ms | 1.15 | 6 | 3.5 | 3.5 | 3.5 |
| **Transfer II** | **HN (dipolar)** | **CAN (dipolar)** | **COCA (scalar)** | **CON (dipolar)** | **COCA (scalar, SCT)** |
| $^{1}$H field / kHz | 95.5 | N/A | N/A | N/A | N/A |
| $^{13}$C field / kHz | N/A | 35.5 | N/A | 27.4 | N/A |
| $^{15}$N field / kHz | 38.1 | 29.2 | N/A | 31.6 | N/A |

| Spectrum | (H)NH | (H)CANH | (H)(CO)CA(CO)NH | (H)CONH | (H)CO(CA)NH |
|---|---|---|---|---|---|
| Carrier $^{13}C$ | N/A | N/A | 173.3 and 53.3 | 173.8 | 173.8 and 53.3 |
| Shape | 100 - 80 % ramp (H) | Tan_ampmod_63_94.land (N) | | Tan_ampmod_63_94.land (N) | |
| Time / ms | 0.85 | 11 | 13.2 | 9 | 6.6 (1st step) 4.6 (2nd step) |
| **Transfer III** | | **NH (dipolar)** | **CON (dipolar)** | **NH (dipolar)** | **CAN (dipolar)** |
| $^{1}H$ field / kHz | N/A | 98.1 | N/A | 98.1 | N/A |
| $^{13}C$ field / kHz | N/A | N/A | 27.4 | N/A | 27.4 |
| $^{15}N$ field / kHz | N/A | 38.6 | 30.3 | 38.1 | 30.3 |
| Carrier $^{13}C$ | N/A | N/A | 173.3 | N/A | 53.3 |
| Shape | N/A | 100 - 80 % ramp (H) | Tan_ampmod_63_94.land (N) | 100 - 80 % ramp (H) | Tan_ampmod_63_94.land (N) |
| Time / ms | N/A | 0.85 | 9 | 0.85 | 9 |
| **Transfer IV** | | | **NH (dipolar)** | | **NH (dipolar)** |
| $^{1}H$ field / kHz | N/A | N/A | 93.7 | N/A | 98.1 |
| $^{13}C$ field / kHz | N/A | N/A | N/A | N/A | N/A |
| $^{15}N$ field / kHz | N/A | N/A | 38.1 | N/A | 38.1 |
| Carrier $^{13}C$ | N/A | N/A | N/A | N/A | N/A |
| Shape | N/A | N/A | 100 - 80 % ramp (H) | N/A | 100 - 80 % ramp (H) |
| Time / ms | N/A | N/A | 0.85 | N/A | 0.85 |
| interscan delay /s | 1 | 1 | 1 | 1 | 1 |

| Spectrum | (H)(CA)CB(CA)NH | (H)(CA)CB(CA)(CO)NH | (H)N(CA)(CO)NH |
|---|---|---|---|
| Scans per point | 12 | 88 | 96 |
| Experimental time / h | 102 | 322.7 | 116 |
| Signal/noise | N/A | N/A | 6.91 |
| Field | 850 | 850 | 850 |
| Spinning frequency/Hz | 55000 | 55000 | 55000 |
| interscan delay /s | 1 | 1 | 1 |
| Sweep width (t1) / ppm | 32(N) | 40(N) | 40.0 (N) |
| Max indirect evolution (t1) / ms | 16.3 (N) | 9.6 (N) | 9.6 (N) |
| Sweep width (t2) / ppm | 90.0 (C) | 84 (C) | 40.0 (N) |

| Spectrum | (H)(CA)CB(CA)NH | (H)(CA)CB(CA)(CO)NH | (H)N(CA)(CO)NH |
|---|---|---|---|
| Max indirect evolution (t2) / ms | 8.8 (C) | 5.6 (C) | 9.6 (N) |
| Sweep width (t3) / ppm | 58.8 (H) | 40.3 (H) | 40.3 (H) |
| Max indirect evolution (t3) / ms | 20.5 (H) | 30.0 (H) | 30.0 (H) |
| **Transfer I** | **HCA (dipolar)** | **HCA (dipolar)** | **HN (dipolar)** |
| $^{1}$H field / kHz | 95.1 | 98.1 | 100 |
| X field / kHz | 40.4 | 40.1 | 38.7 |
| Shape | 85 - 100 % ramp (H) | 85 - 100 % ramp (H) | 100 - 80 % ramp (H) |
| Carrier $^{13}$C | 53.7 | 53.7 | N/A |
| Time / ms | 7 | 3.5 | 1.15 |
| **Transfer II** | **CACBCA(scalar)** | **CACBCA(scalar)** | **NCA (dipolar)** |
| $^{1}$H field / kHz | N/A | N/A | N/A |
| $^{13}$C field / kHz | N/A | N/A | 27.4 |
| $^{15}$N field / kHz | N/A | N/A | 29.4 |
| Carrier $^{13}$C | 53.7 and 48 | 53.7 and 48 | 53.3 |
| Shape | N/A | N/A | Tan_ampmod_63_94.land (N) |
| Time / ms | 12 | 12 | 9 |
| **Transfer III** | **CAN (dipolar)** | **COCA (scalar)** | **CACO (scalar)** |
| $^{1}$H field / kHz | N/A | N/A | N/A |
| $^{13}$C field / kHz | 35.5 | N/A | N/A |
| $^{15}$N field / kHz | 29.2 | N/A | N/A |
| Carrier $^{13}$C | 53.3 | 173.3 and 53.3 | 53.3 and 173.3 |
| Shape | Tan_ampmod_63_94.land (N) | | N/A |
| Time / ms | 11 | 6.6 (1st step) 4.6 (2nd step) | 4.6 (1st step) 6.6 (2nd step) |
| **Transfer IV** | **NH (dipolar)** | **CON (dipolar)** | **CON (dipolar)** |
| $^{1}$H field / kHz | 98.1 | N/A | N/A |
| $^{13}$C field / kHz | N/A | 27.4 | 27.4 |
| $^{15}$N field / kHz | 38.6 | 28.9 | 30.1 |
| Carrier $^{13}$C | N/A | 173.3 | 173.3 |

| Spectrum | (H)(CA)CB(CA)NH | (H)(CA)CB(CA)(CO)NH | (H)N(CA)(CO)NH |
|---|---|---|---|
| Shape | 100 - 80 % ramp (H) | Tan_ampmod_63_94.land (N) | Tan_ampmod_63_94.land (N) |
| Time / ms | 0.85 | 9 | 9 |
| **Transfer V** | | **NH (dipolar)** | **NH (dipolar)** |
| $^{1}$H field / kHz | N/A | 100 | 95.1 |
| $^{13}$C field / kHz | N/A | N/A | N/A |
| $^{15}$N field / kHz | N/A | 38.1 | 38.7 |
| Carrier $^{13}$C | N/A | N/A | N/A |
| Shape | N/A | 100 - 80 % ramp (H) | 100 - 80 % ramp (H) |
| Time / ms | N/A | 0.85 | 0.85 |

| Spectrum | (H)N(CO)(CA)NH | (H)COCA(N)H | H(H)NH |
|---|---|---|---|
| Scans per point | 128 | 72 | 8 |
| Experimental time / h | 154.9 | 158.1 | 22 |
| Signal/noise | 7.85 | 11.9 | 104.84 |
| Field | 850 | 850 | 850 |
| Spinning frequency/Hz | 55000 | 55000 | 55000 |
| interscan delay /s | 1 | 1 | 1 |
| Sweep width (t1) / ppm | 40.0 (N) | 18.0 (CO) | 50.0 (N) |
| Max indirect evolution (t1) / ms | 9.6 (N) | 9.9 (CO) | 12.3 (N) |
| Sweep width (t2) / ppm | 40.0 (N) | 30 (CA) | 15.0 (H) |
| Max indirect evolution (t2) / ms | 9.6 (N) | 8.1 (CA) | 3.5 (H) |
| Sweep width (t3) / ppm | 40.3 (H) | 40.3 (H) | 40.3 (H) |
| Max indirect evolution (t3) / ms | 30.0 (H) | 30.0 (H) | 30.0 (H) |
| **Transfer I** | **HN (dipolar)** | **HCO (dipolar)** | **HH (NOE)** |
| $^{1}$H field / kHz | 100 | 96.1 | N/A |
| X field / kHz | 38.7 | 40.1 | N/A |
| Shape | 100 - 80 % ramp (H) | 85 - 100 % ramp (H) | N/A |
| Carrier $^{13}$C | N/A | 173.3 | N/A |
| Time / ms | 1.15 | 3.5 | 100 |

| Spectrum | (H)N(CO)(CA)NH | (H)COCA(N)H | H(H)NH |
|---|---|---|---|
| **Transfer II** | **NCO(dipolar)** | **COCA (scalar, SCT)** | **HN (dipolar)** |
| $^{1}H$ field / kHz | N/A | N/A | 98.4 |
| $^{13}C$ field / kHz | 27.4 | N/A | N/A |
| $^{15}N$ field / kHz | 30.1 | N/A | 40.5 |
| Carrier $^{13}C$ | 173.3 | 173.3 then 53.3 | N/A |
| Shape | Tan_ampmod_63_94.land (N) | | 80 - 100 % ramp (H) |
| Time / ms | 9 | 6.6 (1st step) 4.6 (2nd step) | **1.15** |
| **Transfer III** | **COCA (scalar)** | **CAN (dipolar)** | **HN (dipolar)** |
| $^{1}H$ field / kHz | N/A | N/A | 95 |
| $^{13}C$ field / kHz | N/A | 27.4 | N/A |
| $^{15}N$ field / kHz | N/A | 27.8 | 40.5 |
| Carrier $^{13}C$ | 173.3 and 53.3 | 53.3 | N/A |
| Shape | N/A | Tan_ampmod_63_94.land (N) | 100 - 80 % ramp (H) |
| Time / ms | 6.6 (1st step) 4.6 (2nd step) | 9 | 0.85 |
| **Transfer IV** | **CAN (dipolar)** | **NH (dipolar)** | |
| $^{1}H$ field / kHz | N/A | 101.9 | N/A |
| $^{13}C$ field / kHz | 27.4 | N/A | N/A |
| $^{15}N$ field / kHz | 29.4 | 38.1 | N/A |
| Carrier $^{13}C$ | 53.3 | N/A | N/A |
| Shape | Tan_ampmod_63_94.land (N) | 100 - 80 % ramp (H) | N/A |
| Time / ms | 9 | 0.85 | N/A |
| **Transfer V** | **NH (dipolar)** | | |
| $^{1}H$ field / kHz | 95.1 | N/A | N/A |
| $^{13}C$ field / kHz | N/A | N/A | N/A |
| $^{15}N$ field / kHz | 38.7 | N/A | N/A |
| Carrier $^{13}C$ | N/A | N/A | N/A |
| Shape | 100 - 80 % ramp (H) | N/A | N/A |
| Time / ms | 0.85 | N/A | N/A |

**Table S4. NMR experimental parameters used in sequence specific assignment in the citrate bound state of.** The powers for soft pulses are taken at the maximum value.

| Spectrum | (H)NH | (H)CANH | (H)(CO)CA(CO)NH | (H)CONH | (H)CO(CA)NH |
|---|---|---|---|---|---|
| Scans per point | 64 | 8 | 56 | 24 | 64 |
| Experimental time / h | 15.9 | 57.6 | 131.4 | 61.9 | 144.9 |
| Signal/noise | 232 | 20.64 | 12.65 | 22.99 | 8.57 |
| Field | 850 | 850 | 850 | 850 | 850 |
| Spinning frequency/Hz | 55000 | 55000 | 55000 | 55000 | 55000 |
| interscan delay /s | 3.5 | 2.5 | 1 | 1 | 0.74 |
| Sweep width (t1) / ppm | 120.0 (N) | 36 (N) | 36 (N) | 34(N) | 34(N) |
| Max indirect evolution (t1) / ms | 24.8 (N) | 17.4 (N) | 15.5 (N) | 18.4 (N) | 18.4 (N) |
| Sweep width (t2) / ppm | 40.8 (H) | 32 (C) | 32 (C) | 16.0 (C) | 16.0 (C) |
| Max indirect evolution (t2) / ms | 29.5 (H) | 7.0 (C) | 6.4 (C) | 12.6 (C) | 12.6 (C) |
| Sweep width (t3) / ppm | N/A | 58.8 (H) | 58.8 (H) | 58.8 (H) | 58.8 (H) |
| Max indirect evolution (t3) / ms | N/A | 20.5 (H) | 20.5 (H) | 20.5 (H) | 20.5 (H) |
| **Transfer I** | **HN (dipolar)** | **HCA (dipolar)** | **HCO (dipolar)** | **HCO (dipolar)** | **HCO (dipolar)** |
| $^{1}$H field / kHz | 96.4 | 94.35 | 93.1 | 91.8 | 94.1 |
| X field / kHz | 70.7 | 40 | 40 | 39.8 | 40.1 |
| Shape | 85 - 100 % ramp (H) | 85 - 100 % ramp (H) | 85 - 100 % ramp (H) | 85 - 100 % ramp (H) | 85 - 100 % ramp (H) |
| Carrier $^{13}$C | | 53.7 | 173.3 | 173.8 | 173.8 |
| Time / ms | 1.2 | 4 | 5 | 4 | 4.2 |
| **Transfer II** | **HN (dipolar)** | **CAN (dipolar)** | **COCA (scalar)** | **CON (dipolar)** | **COCA (scalar)** |
| $^{1}$H field / kHz | 96.4 | N/A | N/A | N/A | N/A |
| $^{13}$C field / kHz | N/A | 34.35 | N/A | 34.4 | N/A |
| $^{15}$N field / kHz | 70.7 | 30.61 | N/A | 31.5 | N/A |
| Carrier $^{13}$C | N/A | 53.7 | 173.3 and 53.3 | 173.8 | 173.8 and 53.3 |

| Spectrum | (H)NH | (H)CANH | (H)(CO)CA(CO)NH | (H)CONH | (H)CO(CA)NH |
|---|---|---|---|---|---|
| Shape | 80 - 100 % ramp (H) | Tan_ampmod_63_94.land (N) | N/A | Tan_ampmod_63_94.land (N) | N/A |
| Time / ms | 1.2 | 13 | 15 | 13 | 7.46 |
| **Transfer III** | | **NH (dipolar)** | **CON (dipolar)** | **NH (dipolar)** | **CAN (dipolar)** |
| $^{1}H$ field / kHz | N/A | 96.45 | N/A | 95.1 | N/A |
| $^{13}C$ field / kHz | N/A | N/A | 34.6 | N/A | 34.9 |
| $^{15}N$ field / kHz | N/A | 39.41 | 32 | 38.7 | 30.1 |
| Carrier $^{13}C$ | N/A | N/A | 173.3 | N/A | 53.3 |
| Shape | N/A | 100 - 80 % ramp (H) | Tan_ampmod_63_94.land (N) | 100 - 80 % ramp (H) | Tan_ampmod_63_94.land (N) |
| Time / ms | N/A | 1.2 | 11.5 | 0.85 | 13 |
| **Transfer IV** | | | **NH (dipolar)** | | **NH (dipolar)** |
| $^{1}H$ field / kHz | N/A | N/A | 96.45 | N/A | 99.2 |
| $^{13}C$ field / kHz | N/A | N/A | N/A | N/A | N/A |
| $^{15}N$ field / kHz | N/A | N/A | 39.41 | N/A | 38.7 |
| Carrier $^{13}C$ | N/A | N/A | N/A | N/A | N/A |
| Shape | N/A | N/A | 100 - 80 % ramp (H) | N/A | 100 - 80 % ramp (H) |
| Time / ms | N/A | N/A | 1.15 | N/A | 0.7 |

| Spectrum | (H)(CA)CB(CA)NH | (H)(CA)CB(CA)(CO)NH | (H)N(CA)(CO)NH |
|---|---|---|---|
| Scans per point | 48 | 48 | 88 |
| Experimental time / h | 296.3 | 269.3 | 262.9 |
| Signal/noise | N/A | N/A | 14.81 |
| Field | 850 | 850 | 800 |
| Spinning frequency/Hz | 55000 | 55000 | 55000 |
| interscan delay /s | 1 | 1 | 1 |
| Sweep width (t1) / ppm | 34(N) | 34(N) | 36.0 (N) |
| Max indirect evolution (t1) / ms | 18.4 (N) | 17.4 (N) | 16.5 (N) |
| Sweep width (t2) / ppm | 70.0 (C) | 70.0 (C) | 36.0 (N) |
| Max indirect evolution (t2) / ms | 6.7 (C) | 6.6 (C) | 16.1 (N) |

| Spectrum | (H)(CA)CB(CA)NH | (H)(CA)CB(CA)(CO)NH | (H)N(CA)(CO)NH |
|---|---|---|---|
| Sweep width (t3) / ppm | 58.8 (H) | 58.8 (H) | 30.1 (H) |
| Max indirect evolution (t3) / ms | 20.5 (H) | 20.5 (H) | 21.3 (H) |
| **Transfer I** | **HCA (dipolar)** | **HCA (dipolar)** | **HN (dipolar)** |
| $^{1}$H field / kHz | 94.7 | 94.4 | 104.2 |
| X field / kHz | 40 | 38.7 | 39.1 |
| Shape | 85 - 100 % ramp (H) | 85 - 100 % ramp (H) | 80 - 100 % ramp (H) |
| Carrier $^{13}$C | 53.7 | 53.7 | N/A |
| Time / ms | 4 | 4 | 1 |
| **Transfer II** | **CACBCA(scalar)** | **CACBCA(scalar)** | **NCA (dipolar)** |
| $^{1}$H field / kHz | N/A | N/A | N/A |
| $^{13}$C field / kHz | N/A | N/A | 20.9 |
| $^{15}$N field / kHz | N/A | N/A | 43.7 |
| Carrier $^{13}$C | 53.7 and 48 | 53.7 and 48 | 53.3 |
| Shape | N/A | N/A | Tan_ampmod_63_94.land (N) |
| Time / ms | 18.56 | 18.56 | 13 |
| **Transfer III** | **CAN (dipolar)** | **COCA (scalar)** | **CACO (scalar)** |
| $^{1}$H field / kHz | N/A | N/A | N/A |
| $^{13}$C field / kHz | 34.8 | N/A | N/A |
| $^{15}$N field / kHz | 30.6 | N/A | N/A |
| Carrier $^{13}$C | 53.7 | 173.3 and 53.3 | 53.3 and 173.3 |
| Shape | Tan_ampmod_63_94.land (N) | | N/A |
| Time / ms | 13 | 7.2 (1st step) 6.4 (2nd step) | 6.4 (1st step) 7.2 (2nd step) |
| **Transfer IV** | **NH (dipolar)** | **CON (dipolar)** | **CON (dipolar)** |
| $^{1}$H field / kHz | 95.4 | N/A | N/A |
| $^{13}$C field / kHz | N/A | 34.4 | 18.4 |
| $^{15}$N field / kHz | 39.4 | 30.6 | 46.5 |
| Carrier $^{13}$C | N/A | 173.3 | 173.3 |
| Shape | 100 - 80 % ramp (H) | Tan_ampmod_63_94.land (N) | Tan_ampmod_63_94.land (N) |

| Spectrum | (H)(CA)CB(CA)NH | (H)(CA)CB(CA)(CO)NH | (H)N(CA)(CO)NH |
|---|---|---|---|
| Time / ms | 1.15 | 13 | 13 |
| **Transfer V** | | **NH (dipolar)** | **NH (dipolar)** |
| $^{1}$H field / kHz | N/A | 96.4 | 95.6 |
| $^{13}$C field / kHz | N/A | N/A | N/A |
| $^{15}$N field / kHz | N/A | 39.4 | 39.1 |
| Carrier $^{13}$C | N/A | N/A | N/A |
| Shape | N/A | 100 - 80 % ramp (H) | 100 - 80 % ramp (H) |
| Time / ms | N/A | 1.15 | 0.6 |

| Spectrum | (H)N(CO)(CA)NH | (H)COCA(N)H | H(H)NH |
|---|---|---|---|
| Scans per point | 144 | 64 | 16 |
| Experimental time / h | 338.6 | 159 | 102.4 |
| Signal/noise | 11.19 | 13.53 | 66.4 |
| Field | 850 | 850 | 850 |
| Spinning frequency/Hz | 55000 | 55000 | 55000 |
| interscan delay /s | 1 | 1 | 1 |
| Sweep width (t1) / ppm | 36.0 (N) | 16.0 (CO) | 60.0 (N) |
| Max indirect evolution (t1) / ms | 14.8 (N) | 12.3 (CO) | 24.8 (N) |
| Sweep width (t2) / ppm | 36.0 (N) | 32 (CA) | 15.0 (H) |
| Max indirect evolution (t2) / ms | 14.8 (N) | 7.6 (CA) | 3.5 (H) |
| Sweep width (t3) / ppm | 58.8 (H) | 58.8 (H) | 40.8 (H) |
| Max indirect evolution (t3) / ms | 20.5 (H) | 20.5 (H) | 29.5 (H) |
| **Transfer I** | **HN (dipolar)** | **HCO (dipolar)** | **HH (NOE)** |
| $^{1}$H field / kHz | 96 | 91.5 | N/A |
| X field / kHz | 38.7 | 39.8 | N/A |
| Shape | 100 - 80 % ramp (H) | 85 - 100 % ramp (H) | N/A |
| Carrier $^{13}$C | N/A | 173.3 | N/A |
| Time / ms | 1.15 | 4 | 100 |
| **Transfer II** | **NCO(dipolar)** | **COCA (scalar, SCT)** | **HN (dipolar)** |

| Spectrum | (H)N(CO)(CA)NH | (H)COCA(N)H | H(H)NH |
|---|---|---|---|
| $^{1}H$ field / kHz | N/A | N/A | 93.6 |
| $^{13}C$ field / kHz | 27.1 | N/A | N/A |
| $^{15}N$ field / kHz | 37.7 | N/A | 38.7 |
| Carrier $^{13}C$ | 173.3 | 173.3 then 53.3 | N/A |
| Shape | Tan_ampmod_63_94.land (N) | | 100 - 80 % ramp (H) |
| Time / ms | 13 | 7.2 (1st step) 6.4 (2nd step) | 0.7 |
| **Transfer III** | **COCA (scalar)** | **CAN (dipolar)** | **HN (dipolar)** |
| $^{1}H$ field / kHz | N/A | N/A | 94.2 |
| $^{13}C$ field / kHz | N/A | 35 | N/A |
| $^{15}N$ field / kHz | N/A | 29.7 | 38.7 |
| Carrier $^{13}C$ | 173.3 and 53.3 | N/A | N/A |
| Shape | N/A | Tan_ampmod_63_94.l and (N) | 100 - 80 % ramp (H) |
| Time / ms | 7.2 (1st step) 6.4 (2nd step) | 12.5 | 0.6 |
| **Transfer IV** | **CAN (dipolar)** | **NH (dipolar)** | |
| $^{1}H$ field / kHz | N/A | 95.1 | N/A |
| $^{13}C$ field / kHz | 35.1 | N/A | N/A |
| $^{15}N$ field / kHz | 30.1 | 38.7 | N/A |
| Carrier $^{13}C$ | 53.3 | N/A | N/A |
| Shape | Tan_ampmod_63_94.land (N) | 100 - 80 % ramp (H) | N/A |
| Time / ms | 12.5 | 0.85 | N/A |
| **Transfer V** | **NH (dipolar)** | | |
| $^{1}H$ field / kHz | 95.1 | N/A | N/A |
| $^{13}C$ field / kHz | N/A | N/A | N/A |
| $^{15}N$ field / kHz | 38.7 | N/A | N/A |
| Carrier $^{13}C$ | N/A | N/A | N/A |
| Shape | 100 - 80 % ramp (H) | N/A | N/A |
| Time / ms | 0.85 | N/A | N/A |

**Table S5. NMR experimental parameters used in sequence specific assignment in the citrate free state of CitApc with IVFL reverse labeling.** The powers for soft pulses are taken at the maximum value.

| Spectrum | (H)NH | (H)CANH | (H)(CO)CA(CO)NH |
|---|---|---|---|
| Scans per point | 136 | 24 | 32 |
| Experimental time / h | 15.1 | 58.8 | 117.3 |
| Signal/noise | 226.17 | 21.43 | N/A |
| Field | 850 | 850 | 850 |
| Spinning frequency/Hz | 55000 | 55000 | 55000 |
| interscan delay /s | 1 | 1 | 1 |
| Sweep width (t1) / ppm | 160 (N) | 40 (N) | 40(N) |
| Max indirect evolution (t1) / ms | 14.5 (N) | 11.0 (N) | 9.6 (N) |
| Sweep width (t2) / ppm | 40.3 (H) | 30 (C) | 84 (C) |
| Max indirect evolution (t2) / ms | 30.0 (H) | 9.0 (C) | 5.6 (C) |
| Sweep width (t3) / ppm | N/A | 40.3 (H) | 40.3 (H) |
| Max indirect evolution (t3) / ms | N/A | 30.0 (H) | 30.0 (H) |
| **Transfer I** | **HN (dipolar)** | **HCA (dipolar)** | **HCA (dipolar)** |
| $^{1}$H field / kHz | 98.2 | 100.9 | 103.6 |
| X field / kHz | 38.1 | 40.1 | 40.1 |
| Shape | 100 - 80 % ramp (H) | 85 - 100 % ramp (H) | 85 - 100 % ramp (H) |
| Carrier $^{13}$C | N/A | 53.7 | 53.7 |
| Time / ms | 1.15 | 3.5 | 3.5 |
| **Transfer II** | **HN (dipolar)** | **CAN (dipolar)** | **CACBCA(scalar)** |
| $^{1}$H field / kHz | 94.5 | N/A | N/A |
| $^{13}$C field / kHz | N/A | 27.4 | N/A |
| $^{15}$N field / kHz | 38.1 | 26.6 | N/A |
| Carrier $^{13}$C | N/A | 53.7 | 53.7 and 48 |
| Shape | 100 - 80 % ramp (H) | Tan_ampmod_63_94.land (N) | |
| Time / ms | 0.8 | 9 | 6 |
| **Transfer III** | | **NH (dipolar)** | **CAN (dipolar)** |
| $^{1}$H field / kHz | N/A | 98.1 | N/A |

| Spectrum | (H)NH | (H)CANH | (H)(CO)CA(CO)NH |
|---|---|---|---|
| $^{13}C$ field / kHz | N/A | N/A | 27.4 |
| $^{15}N$ field / kHz | N/A | 38.1 | 26.6 |
| Carrier $^{13}C$ | N/A | N/A | 53.3 |
| Shape | N/A | 100 - 80 % ramp (H) | Tan_ampmod_63_94.land (N) |
| Time / ms | N/A | 0.85 | 9 |
| **Transfer IV** | | | **NH (dipolar)** |
| $^{1}H$ field / kHz | N/A | N/A | 97.2 |
| $^{13}C$ field / kHz | | N/A | N/A |
| $^{15}N$ field / kHz | N/A | N/A | 38.1 |
| Carrier $^{13}C$ | N/A | N/A | N/A |
| Shape | N/A | N/A | 100 - 80 % ramp (H) |
| Time / ms | N/A | N/A | 0.85 |

**Table S6. NMR experimental parameters used in sequence specific assignment in the citrate bound state of CitApc with IVFL reverse labeling.** The powers for soft pulses are taken at the maximum value.

| Spectrum | (H)NH | (H)CANH | (H)(CO)CA(CO)NH | (H)(CA)CB(CA)NH | (H)(CA)CB(CA)(CO)NH |
|---|---|---|---|---|---|
| Scans per point | 80 | 16 | 64 | 32 | 72 |
| Experimental time / h | 6.2 | 85.3 | 153.5 | 158.4 | 337.9 |
| Signal/noise | 181.46 | 21.65 | 10.86 | N/A | N/A |
| Field | 850 | 850 | 850 | 850 | 850 |
| Spinning frequency/Hz | 55000 | 55000 | 55000 | 55000 | 55000 |
| interscan delay /s | 1 | 2 | 1 | 1 | 1 |
| Sweep width (t1) / ppm | 120 (N) | 36 (N) | 36 (N) | 36(N) | 34(N) |
| Max indirect evolution (t1) / ms | 13.5 (N) | 16.1 (N) | 14.5 (N) | 14.5 (N) | 15.0 (N) |
| Sweep width (t2) / ppm | 40.8 (H) | 32 (C) | 32 (C) | 70.0 (C) | 70 (C) |
| Max indirect evolution (t2) / ms | 29.5 (H) | 7.0 (C) | 7.0 (C) | 6.6 (C) | 6.4 (C) |
| Sweep width (t3) / ppm | N/A | 58.8 (H) | 58.8 (H) | 58.8 (H) | 58.8 (H) |
| Max indirect evolution (t3) / ms | N/A | 20.5 (H) | 20.5 (H) | 20.5 (H) | 20.5 (H) |
| **Transfer I** | **HN (dipolar)** | **HCA (dipolar)** | **HCO (dipolar)** | **HCA (dipolar)** | **HCA (dipolar)** |
| $^{1}H$ field / kHz | 96 | 92.9 | 82.6 | 90.7 | 87.3 |
| X field / kHz | 38.1 | 41 | 41 | 41 | 37.7 |
| Shape | 100 - 80 % ramp (H) | 85 - 100 % ramp (H) | 85 - 100 % ramp (H) | 85 - 100 % ramp (H) | 85 - 100 % ramp (H) |
| Carrier $^{13}C$ | | 53.7 | 173.3 | 53.7 | 53.7 |
| Time / ms | 1 | 6 | 6 | 5.5 | 4 |
| **Transfer II** | **HN (dipolar)** | **CAN (dipolar)** | **COCA (scalar)** | **CACBCA(scalar)** | **CACBCA(scalar)** |
| $^{1}H$ field / kHz | 95.4 | N/A | N/A | N/A | N/A |
| $^{13}C$ field / kHz | N/A | 36 | N/A | N/A | N/A |

| **Spectrum** | (H)NH | (H)CANH | (H)(CO)CA(CO)NH | (H)(CA)CB(CA)NH | (H)(CA)CB(CA)(CO)NH |
|---|---|---|---|---|---|
| $^{15}$N field / kHz | 38.1 | 28.4 | N/A | N/A | N/A |
| Carrier $^{13}$C | N/A | 53.7 | 173.3 and 53.3 | 53.7 and 48 | 53.7 and 48 |
| Shape | 100 - 80 % ramp (H) | Tan_ampmod_63_94.land (N) | | N/A | N/A |
| Time / ms | 0.8 | 11 | 15.6 | 18.4 | 18.4 |
| **Transfer III** | | **NH (dipolar)** | **CON (dipolar)** | **CAN (dipolar)** | **COCA (scalar)** |
| $^{1}$H field / kHz | N/A | 95.4 | N/A | N/A | N/A |
| $^{13}$C field / kHz | N/A | N/A | 35.4 | 36 | N/A |
| $^{15}$N field / kHz | N/A | 38.1 | 31 | 28.9 | N/A |
| Carrier $^{13}$C | N/A | N/A | 173.3 | 53.3 | 173.3 and 53.3 |
| Shape | N/A | 100 - 80 % ramp (H) | Tan_ampmod_63_94.land (N) | Tan_ampmod_63_94.land (N) | |
| Time / ms | N/A | 1.15 | 11 | 11 | 15.6 |
| **Transfer IV** | | | **NH (dipolar)** | **NH (dipolar)** | **CON (dipolar)** |
| $^{1}$H field / kHz | N/A | N/A | 92.2 | 89.6 | N/A |
| $^{13}$C field / kHz | N/A | N/A | N/A | N/A | 33.5 |
| $^{15}$N field / kHz | N/A | N/A | 38.1 | 38.1 | 30.3 |
| Carrier $^{13}$C | N/A | N/A | N/A | N/A | 173.3 |
| Shape | N/A | N/A | 100 - 80 % ramp (H) | 100 - 80 % ramp (H) | Tan_ampmod_63_94.land (N) |
| Time / ms | N/A | N/A | 1.15 | 0.8 | 13 |
| **Transfer V** | | | | | **NH (dipolar)** |
| $^{1}$H field / kHz | N/A | N/A | N/A | N/A | 88.7 |
| $^{13}$C field / kHz | N/A | N/A | N/A | N/A | N/A |
| $^{15}$N field / kHz | N/A | N/A | N/A | N/A | 38.1 |
| Carrier $^{13}$C | N/A | N/A | N/A | N/A | N/A |
| Shape | N/A | N/A | N/A | N/A | 100 - 80 % ramp (H) |
| Time / ms | N/A | N/A | 0.85 | N/A | 0.85 |

**Table S7. Parameters extracted from CEST profile fitting of the WT PASc domain.**

| | exchange rates | | | | |
|---|---|---|---|---|---|
| Kab [s-1] | 17.731 +/- 5.147 | | | | |
| Kba [s-1] | 450.471 +/- 94.875 | | | | |
| residue | Peak position [ppm] | CS difference [ppm] | R1 [s-1] | R2a [s-1] | R2b [s-1] |
| E209 | 116.9445 +/- 0.0709 | 3.5118 +/- 0.2071 | 1.1831 +/- 0.0213 | 87.1442 +/- 6.2674 | 38.4808 +/- 75.2772 |
| E215 | 114.7406 +/- 0.0415 | 4.5057 +/- 0.2311 | 0.4289 +/- 0.0215 | 62.1840 +/- 4.4704 | 74.8535 +/- 79.2006 |

**Table S8. Parameters extracted from CEST profile fitting of the N288D mutant PASc domain.**

| | exchange rates | | | | |
|---|---|---|---|---|---|
| Kab [s-1] | 2.409 +/- 0.281 | | | | |
| Kba [s-1] | 70.442 +/- 13.434 | | | | |
| residue | Peak position [ppm] | CS difference [ppm] | R1 [s-1] | R2a [s-1] | R2b [s-1] |
| A212 | 120.4246 +/- 0.0549 | 2.8159 +/- 0.1641 | 0.4981 +/- 0.0163 | 34.2863 +/- 2.2802 | 38.8838 +/- 56.3272 |
| L214 | 117.8093 +/- 0.0378 | 5.2163 +/- 0.1215 | 0.3986 +/- 0.0157 | 31.4040 +/- 2.0766 | 142.6128 +/- 33.4526 |
| E215 | 113.6648 +/- 0.0406 | 5.2638 +/- 0.1194 | 0.5784 +/- 0.0218 | 36.4141 +/- 2.4877 | 254.6045 +/- 60.6037 |
| I217 | 114.0103 +/- 0.0362 | 4.9695 +/- 0.1101 | 0.5383 +/- 0.0147 | 36.4148 +/- 2.6105 | 126.8150 +/- 24.4825 |

**Table S9. X-ray data collection statistics GT PASp-Citrate complex.**

| | |
|---|---|
| Wavelength | 0.99979 Å |
| Beamline | SLS-X10SA |
| Detector | PILATUS 6M |
| Space group | $P2_1$ |
| *a* | 64.21 Å |
| *b* | 101.14 Å |
| *c* | 77.88 Å |
| α, γ | 90° |
| *β* | 105.63° |
| MOL (AU) | 8 |
| Resolution[a] | 48.79-1.70 Å (1.73-1.70 Å) |
| No. unique reflections | 105,114 |
| Redundancy | 6.85 (6.86) |
| Completeness(%) | 99.9 (99.5) |
| Mean $I/\sigma(I)$ | 16.47 (2.71) |
| $R_{means}$ (%) | 4.85 (29.52) |

[a]Values in parentheses are outer-resolution shell.

**Table S10. X-ray data collection statistics GT PASp-R93A.**

| | |
|---|---|
| Wavelength | 0.99979 Å |
| Beamline | SLS-X10SA |
| Detector | PILATUS 6M |
| Space group | $C222_1$ |
| *a* | 60.14 Å |
| *b* | 120.05 Å |
| *c* | 152.14 Å |
| α, *β*, γ | 90° |
| MOL (AU) | 4 |
| Resolution[a] | 47.12-1.61 Å (1.64-1.61 Å) |
| No. unique reflections | 70,580 |
| Redundancy | 12.94 (10.44) |
| Completeness(%) | 99.1 (93.3) |
| Mean *I*/σ (*I*) | 16.33 (1.27) |
| $R_{meas}$ (%) | 4.89 (63) |

[a]Values in parentheses are outer-resolution shell.

**Table S11. X-ray data collection and phasing statistics GT PASc-N288D.**

| | Peak (PEAK) | Inflection (INFL) | High-energy remote (HREM) |
|---|---|---|---|
| Data collection statistics | | | |
| Wavelength (Å) | 0.97957 | 0.9800 | 0.97188 |
| X-ray source | | X10SA, Swiss light source | |
| Detector | | PILATUS 6M | |
| Space group | | $P2_12_12_1$ | |
| Unit cell parameters[a] | *a*=48.82 Å, *b*=49.42 Å, *c*=91.77 Å, α=β=γ=90° | *a*=49.31 Å, *b*=49.38 Å, *c*=92.92 Å, α=β=γ=90° | *a*=48.94 Å, *b*=49.31 Å, *c*=92.19 Å, α=β=γ=90° |
| MOL (AU) | | 2 | |
| Resolution (Å) | 43.51-2.13 (2.17-2.13) | 46.35-2.50 (2.56-2.50) | 43.51-2.10 (2.15-2.10) |
| No. unique reflections | 12,062 | 8245 | 12,630 |
| Redundancy | 3.17 (2.60) | 12.84 (13.39) | 3.13 (2.51) |
| Completeness (%) | 93.1 (83.9) | 99.5 (100) | 92.5 (80.1) |
| Mean I/(σI) | 15.75 (2.76) | 21.26 (3.67) | 20.96 (3.0) |
| $R_{meas}$ (%) | 5.08 (28.98) | 3.76 (21.80) | 3.82 (26.67) |
| Substructure solution (SHELX) | | | |
| d´´/sig | 1.10 at 3.28-2.98 Å | 1.24 at 5.52-4.38 Å | 1.13 at 3.71-3.24 Å |
| $CC_{1/2}$ | 85.1 (Inf - 4.74 Å) | 87.7 (Inf - 5.52 Å) | 86.9 (Inf - 4.67 Å) |
| $CC_{HREM/PEAK}$ | | 44.1 % at 3.5 - 3.1 Å | |
| $CC_{HREM/INFL}$ | | 34.9 % at 3.5 - 3.1 Å | |
| $CC_{PEAK/INFL}$ | | 37.8 % at 3.5 - 3.1 Å | |
| No. of sites | | 4 | |
| $CC_{ALL/WEAK}$ | | 44.6/29.6 | |

[a]Values in parentheses are outer-resolution shells

**Table S12. X-ray structure refinement statistics GT PASp-Citrate complex.**

| | |
|---|---|
| R-factor | 16.7 |
| $R_{free}$[a] | 19.6 |
| Solvent (%) | 43.52 |
| Mean B-value (Å$^2$) | |
| Protein (main/side) | 23.0/28.54 |
| citrate | 20.62 |
| sodium | 33.29 |
| water | 35.9 |
| No. of protein residues | 1029 |
| No. of water residues | 726 |
| No. of citrate molecules | 8 |
| No. of sodium atoms | 1 |
| Root mean square deviations from ideal geometry | |
| Bond lengths (Å) | 0.011 |
| Bond angles (°) | 1.65 |
| Ramachandran plot (%) | |
| Favoured | 97.84 |
| Allowed | 2.16 |
| Outliers | 0 |

[a]$R_{free}$ was determined using 4.956 % of the data (*60*)

**Table S13. X-ray structure refinement statistics GT PASp-R93A.**

| | |
|---|---|
| R-factor | 19.1 |
| $R_{free}$[a] | 22.3 |
| Solvent (%) | 50.25 |
| Mean B-value (Å$^2$) | |
| Protein (main/side) | 31.38/36.52 |
| CXS[b] | 47.69 |
| sulfate | 43.24 |
| glycerol | 44.66 |
| water | 41.56 |
| No. of protein residues | 500 |
| No. of water residues | 221 |
| No. of CXS molecules | 4 |
| No. of sulfate atoms | 3 |
| No. of glycerol molecules | 2 |
| r.m.s.d. | |
| Bond lengths (Å) | 0.007 |
| Bond angles (°) | 0.975 |
| Ramachandran plot (%) | |
| Favoured | 97.02 |
| Allowed | 2.73 |
| Outliers | 0.25 |

[a]$R_{free}$ was determined using 5.08 % of the data (*60*)
[b]3-Cyclohexyl-1-propylsulfonic acid

**Table S14. X-ray structure refinement statistics GT PASc-N288D.**

| | |
|---|---|
| R-factor | 19.5 |
| $R_{free}$[a] | 23.3 |
| Solvent (%) | 46.31 |
| Mean B-value (Å$^2$) | |
| Protein (main/side) | 43.25/50.91 |
| water | 42.53 |
| Mg | 50.77 |
| No. of protein residues | 219 |
| No. of water residues | 33 |
| No. of magnesium atoms | 1 |
| r.m.s.d. | |
| Bond lengths (Å) | 0.009 |
| Bond angles (°) | 1.614 |
| Ramachandran plot (%) | |
| Favoured | 96.17 |
| Allowed | 3.83 |
| Outliers | 0 |

[a]$R_{free}$ was determined using 5.067 % of the data (*60*)
